# Life-cycle trajectory inference links temperature-gated progenitors to reproductive fate

**DOI:** 10.64898/2026.06.29.735214

**Authors:** Gaurav Vaidya, Elena Lagodny, Aishwarya Girish, Ana Maria Mellado Fuentes, Eric Ross, Sofia Robb, Kristina Mirkes, Deniz Yavru, Arif Ul Maula Khan, Christian Tischer, Michael W. Dorrity, Hanh Thi-Kim Vu

## Abstract

Temperature shapes reproductive strategies across animals, yet how individuals switch between sexual and asexual reproduction remains unknown. We establish the planarian *Phagocata morgani* as a model for temperature-dependent reproductive plasticity and adapt multiplexed single-cell transcriptomics to profile >1 million nuclei from >300 animals across body sizes and temperatures. Leveraging individual variation in cell composition, we reconstruct an organism-wide trajectory that bifurcates toward alternative reproductive fates. Temperature extremes constrain worms to one fate, whereas intermediate conditions permit probabilistic commitment to either. At the bifurcation, temperature gates a stem cell pool: warmth suppresses differentiation and promotes progenitor accumulation, whereas cold transcriptionally activates this pool for *de novo* sexual organogenesis. These findings reveal how environmental inputs act on stem cells to couple body size, temperature, and reproductive fate.

## Introduction

Environmental temperature shapes animal development, physiology, and reproductive mode (*1*, *2*). One striking example is facultative reproduction, in which a single organism adopts a sexual or asexual strategy depending on intrinsic state and environmental conditions (*3–5*). Sexual reproduction promotes genetic diversity and can increase fitness in fluctuating environments, whereas asexual reproduction allows rapid population expansion when mates are absent or conditions favor clonal propagation (*6*, *7*). Each mode also carries costs: sexual reproduction requires investment in finding mates and developing reproductive tissues, while asexual reproduction limits genetic variation among offspring (*8*, *9*). Facultative reproduction strategically links reproductive mode to environmental context, yet the cellular mechanisms allowing an individual animal to switch between sexual and asexual fates are unknown, in part due the rarity of switching among classical animal models.

Planarian flatworms are uniquely suited to this problem because they combine extensive adult developmental plasticity with remarkable reproductive diversity. In particular, *Phagocata morgani* (Fig. 1A), a freshwater planarian native to eastern North America, reproduces sexually through *de novo* production of a hermaphroditic reproductive system in winter, but propagates by asexually fissioning through the warmer months (*10*) (Fig. 1B). This adult developmental plasticity relies on neoblasts, pluripotent stem cells that form all somatic and potentially germline lineages, and continuously maintain whole-body turnover of all tissues and organs (*11–13*). Thus, a change in reproductive mode must be integrated with ongoing growth, tissue maintenance, and organogenesis across the entire adult body. This raises a central question: how does one animal generate two distinct reproductive phenotypes from the same cellular system, and how does temperature bias this decision?

**Fig. 1.**
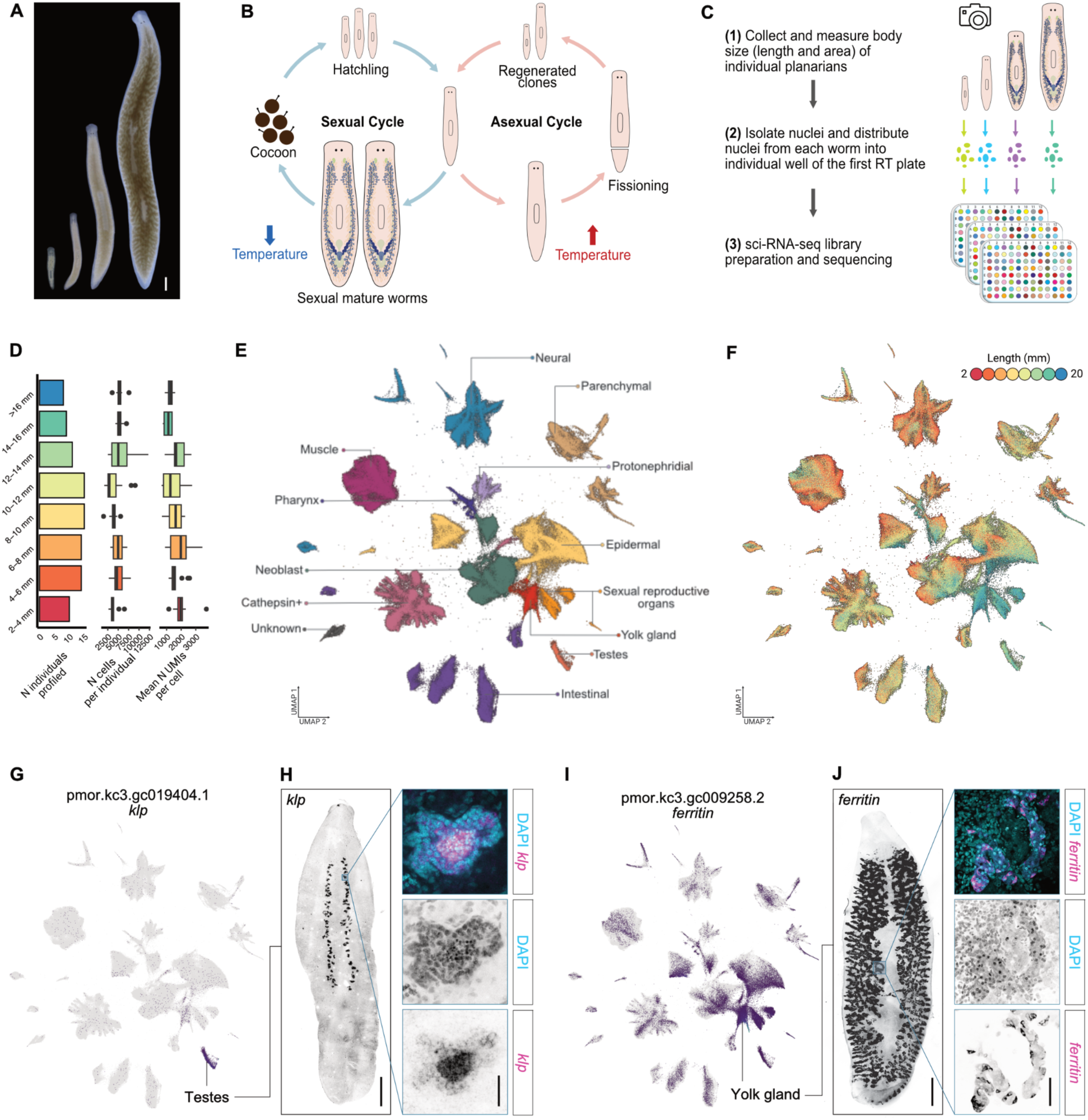
An individual-resolved single-cell atlas of *P. morgani* across body size. (**A**) Live images of *P. morgani* at different body sizes. Scale bar: 1 mm. (**B**) Life-cycle schematic detailing how P. morgani alternates between sexual and asexual reproduction (binary fission) in response to temperature. (**C**) Experimental workflow for generating the individually resolved single-cell atlas of P. morgani across body sizes. (**D**) Number of worms profiled (left), cells profiled per individual worm (center), and mean UMI counts per cell per worm (right). (**E**) Two-dimensional uniform manifold approximation and projection (UMAP) embedding, colored by broad cell type annotation. (**F**) Two-dimensional UMAP embedding, colored by individual worm length. (**G**) UMAP embedding colored by expression of *klp* (pmor.kc3.gc019404.1), a marker of the testes. (**H**) Whole-mount FISH of *klp* (left), scale bar: 1 mm; insets show a high-magnification view of the testes with klp (magenta) and DAPI (cyan) merged (top), DAPI alone (middle), and *klp* alone (bottom), scale bar: 50 μm. (I) UMAP embedding colored by expression of ferritin (pmor.kc3.gc009258.2), a marker of the yolk gland. (J) Whole-mount FISH of ferritin (left), scale bar: 1 mm; insets show a high-magnification view of the yolk gland with ferritin (magenta) and DAPI (cyan) merged (top), DAPI alone (middle), and ferritin alone (bottom), scale bar: 50 μm.

Resolving this question requires connecting environmental input to organism-wide cellular change. Temperature could act (i) globally by altering organismal growth and physiology or (ii) locally via a subset of cells to drive the animal toward sexual or asexual fate. Distinguishing between these possibilities requires cellular measurements across many individuals differing in body size, temperature exposure and reproductive state. Single-cell RNA sequencing provides the necessary molecular and cellular resolution, but standard approaches that pool cells across individuals obscure the animal-to-animal variation needed to disentangle organism-level decisions. Capturing cellular variation across animals would allow reconstruction of organism-level developmental trajectories, akin to the cellular trajectories used to disentangle fate decisions. Because biological systems with many degrees of freedom often vary along fewer effective dimensions (*14–17*), individual-resolved single-cell profiling can use variation among animals or samples to infer latent axes such as developmental progression, maturation, and disease state (*14*, *16*). In *P. morgani*, where the reproductive state emerges in the context of continuous body growth, preserving individual identity is essential to separate size-dependent remodeling from temperature-dependent bias in reproductive fate.

Here, we establish *P. morgani* as a tractable model system for studying temperature-dependent reproductive plasticity. We adapted sci-RNA-seq3 (*18*), a multiplexed single-cell combinatorial indexing approach, to *P. morgani* and generated an individually resolved single-cell atlas comprising over one million nuclei from more than 300 worms spanning a broad range of body sizes and temperatures. By tracking shifts in cell composition across individuals, we computationally reconstructed the growth trajectory underlying this facultative reproductive decision. We found that the growth trajectory of *P. morgani* undergoes a sharp developmental bifurcation shaped by body size and gated by temperature. One branch leads to a sexual state, marked by the emergence of testes, yolk gland cells, and other reproductive cell types. The other branch is asexually biased and is defined by an enrichment of subpopulations of epidermal, pharynx and parenchymal cells, as well as an absence of reproductive cells. Using statistical analysis of cell-cell growth covariation throughout the body, we pinpointed a single neoblast subpopulation, Neoblast 5, positioned at the intersection of the two branches of reproductive fate. Profiling animals grown across a controlled temperature gradient revealed that the abundance and activation of this pool are decoupled. Under warm, asexually promoting conditions, Neoblast 5 cells accumulate while sexual lineages are depleted. Under cold, sexually promoting conditions, Neoblast 5 cells become transcriptionally active and are depleted as reproductive tissues emerge. Together, these results suggest that temperature acts not by broadly altering growth or physiology, but through a specific cold-activated neoblast state that links body size and environmental context to reproductive fate, providing a cellular basis for phenotypic plasticity.

## Results

### An individual-resolved single-cell atlas of *P. morgani* across body size

Body size plays a critical role in determining the onset of reproduction in both sexual (*19*) and asexual strains of the model planarian species *S. mediterranea* (*20*). To investigate facultative reproduction in the non-model planarian *P. morgani*, we first established husbandry methods and generated resources for genomic analysis, including a high-quality transcriptome assembly and gene annotation (Materials and Methods). With these resources in place, we set out to generate the first comprehensive single-cell atlas for this species. To capture reproductive modes of *P. morgani*, we sampled 96 worms ranging from 2-18 mm (0.158 mm resolution on average) and performed sci-RNA-seq3 (Fig. 1D). We adapted the published sci-RNA-seq3 protocol (*18*, *21*) to *P. morgani*, introducing modifications that increased reverse transcription efficiency and reduced erroneous combinatorial barcodes (Fig. 1C; Materials and Methods). We measured body size for each individual worm captured in the sci-RNA-seq3 experiment using a custom image analysis pipeline capable of extracting length and area from short video captures (Fig. S1A-B). After quality control filtering (Fig. S1C), we captured a total of 446,653 single-nuclei transcriptomes (median 1,240 UMIs and 774 genes detected per cell) with a per-worm recovery range of 1,400-12,000 nuclei across the 96 individuals (Fig. 1D). We grouped cells into 56 clusters and annotated cell type identity using marker gene expression per cluster (Table S1) and comparison to previously published single-cell datasets for the model planarian *S. mediterranea* (*22*, *23*). We identified a total of 14 broad cell types, 55 cell types and 101 sub cell types (Fig. 1E; Fig. S1D; Table S2-S3). For this level of cellular complexity, a median of 4,344 cells recovered per worm provides ∼40x cellular coverage, enabling high-resolution analysis of growth dynamics for all cell types in *P. morgani*.

Strikingly, several cell type clusters showed strong size specificity, composed predominantly of cells from worms larger than 12 mm in length (Fig. 1F). Using sexual reproductive system markers in the model species *S. mediterranea*, such as kinesin-like protein-14E and ferritin (*24–26*), we identified these size-specific cell types as components of the reproductive system (Fig. 1G, I; Fig. S1E), consistent with the onset of sexual maturation at larger body sizes. We validated these reproductive cell types by fluorescence in situ hybridization (FISH) using cluster-specific markers (Fig. 1H, J; Fig. S1J) and, in addition, validated a few other cell types typical of freshwater planarians (Fig. S1E-H). Similar to the distribution in the model *S. mediterranea*, the testes run parallel to the ventral nerve cords, however, in *P. morgani* they do not span the entire length of the body (*24*) (Fig. 1H). The yolk gland on the other hand matches the distribution pattern observed in *S. mediterranea* (*18*) (Fig. 1J). Overall, we establish a new planarian model system, characterize its entire cellular composition, and identify the *de novo* development of the reproductive apparatus along its growth trajectory.

### Analysis of cell composition resolves a forked growth trajectory in *P. morgani*

To understand the relationship between body size and cell composition, we extracted cell type counts per sample to build a worm-by-cell-type matrix (Fig. 2A) and examined how cell type proportions changed as a function of body size. We observed that every cell type in the worm showed size dependence, reflecting the profound reorganization of cell proportions during growth (Fig. 2B; Fig. S2A). For example, neoblasts, the pluripotent stem cells in planarians, increased in abundance with body size, while other cell types like muscle or neural decreased with size (Fig. 2B). To understand the collective variation in cell growth dynamics across the worm, we used Principal Component Analysis (PCA) to visualize cell composition in three dimensions, with each point representing an individual worm and its position on the dominant axes (72.5% total variance explained in PCs1-3) of cellular variation (*14*, *27*) (Fig. 2C). We observed that points were ordered in this space roughly according to body size (Fig. 2C), consistent with the strong size-dependent remodeling of cell abundances we observed previously. We then constructed a principal graph to infer a growth trajectory through these points (*20*, *27*) (Fig. 2C). The trajectory originated from the smallest worms and tracked with body size until ∼8 mm, where an unexpected branching event separated the largest worms in the dataset from a smaller set of worms ∼8-11 mm in size (Fig. 2C).

**Fig. 2.**
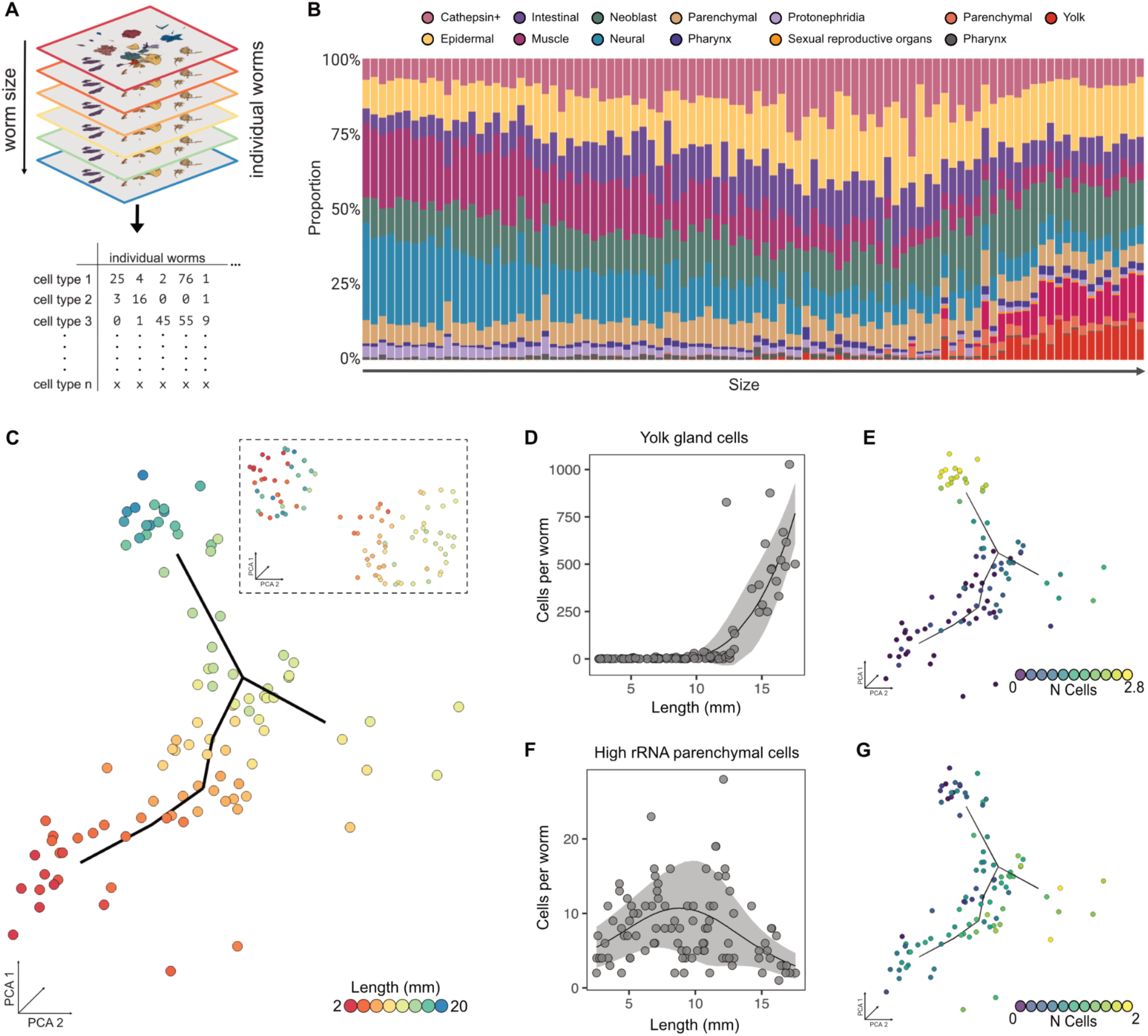
Reconstruction of the facultative sexual trajectory of P. morgani from whole-organism cell composition. (**A**) Schematic of how per-cell gene expression matrices are aggregated into a per-worm cell-composition matrix (cell types × individual worms). (**B**) Cell-composition dynamics during growth: the proportion of each broad cell type per worm, with worms arranged in ascending order of body length (left to right). (**C**) Growth trajectory inferred by pseudotime ordering (Monocle3) through points in PCA space; each point is a single worm colored by body length. Inset: the same worms after regressing out the size effect (size-corrected PCA). (**D**) Yolk gland cell abundance (cells per worm) across the sampled size range; line is the fitted trend and the ribbon indicates the standard deviation. (**E**) The same cell type plotted along the growth trajectory, with each worm colored by its normalized cell count (log₁₀). (**F**) High rRNA parenchymal cell abundance across the sampled size range, plotted as in (D). (**G**) High rRNA parenchymal cells along the growth trajectory, colored as in (E).

To determine if the branches reflected the two reproductive modes of the worm, we analyzed cell abundances that drive the trajectory’s structure. We first analyzed relative abundances of known cell types associated with sexual reproduction such as testes cells and yolk gland cells (*24*, *28–30*). Plotting these abundances (Fig. 2D-E; Fig. S2B-C) in the PCA space showed that they align with the branch containing the largest worms, suggesting that the worms at the end of this branch are committed to reproducing sexually. Without any known asexual-associated cell types from other planarians, we performed an unsupervised characterization of the second branch. We clustered the worms in the PCA space and identified four distinct groups which were annotated based on their position on the inferred trajectory (Fig. S2D). We identified cell-types whose abundances were enriched at each position on the trajectory, including those defining the putative asexual reproductive fate cluster (Table S4; Fig. 2F-G; Fig. S2E-F). This branch was defined by worms of intermediate size that lack the reproductive cell signature of the opposing branch, and were enriched for subpopulations of epidermal, pharynx and parenchymal cells amongst others.

To assess whether the branching trajectory reflects distinct reproductive fates or if its structure reflects size variation in worms that differ in cell composition by chance, we analyzed size-independent variation. We removed size effects by regressing each PC on body size and subtracted the fitted size component from each PC, generating a corrected PCA space to visualize any residual variation in cell composition among worms (Materials and Methods). Reducing dimensions of this corrected PCA space using UMAP, we found that the bifurcation persisted, with sexual and asexual branches forming two distinct clusters occupied by worms of varying sizes (inset of Fig. 2C). To verify that these clusters retained their specific reproductive identities rather than reflecting an unrelated axis of variation, we visualized relative abundances of branch-defining cell-type markers in the size-corrected UMAP space (Fig. S2G-H). Sexual reproductive markers remained sharply concentrated within one cluster and were virtually absent from the other. Conversely, markers associated with the asexual branch exhibited the inverse distribution, recapitulating the branch identities from our initial analysis. Thus, cell composition variation in the worm is pulled along separable axes of variation: (i) body size and (ii) reproductive mode. The complex remodeling of cells in the animal can be reduced to a simple branching geometry inferred from dozens of individual worms. Taken together, these results suggest that *P. morgani* commits to either asexual or sexual trajectories at a specific point during growth, with worms committed to sexual reproduction ultimately reaching larger body sizes.

### Uncovering cellular drivers of a whole-animal reproductive fate decision

We next sought to identify cell types whose abundance commits the worm to one reproductive fate over the other, reasoning that a cellular trigger for *de novo* sexual organ production would (i) show significant covariation with other cell types in the worm and (ii) act independently of size-dependent cellular remodeling. To pinpoint cellular drivers with this behavior, we modeled covariance structure across cell types using Poisson lognormal networks (PLNmodels) implemented in Hooke, using worm body size as a covariate to isolate size-independent effects (*31*, *32*). Cell-cell covariance contains latent information on developmental relationships - for example, a progenitor population is expected to show negative covariation with its descendants while sister lineages with shared growth dynamics are expected to show positive covariation (Fig. 3A). Because commitment to sexual fate requires *de novo* production of the reproductive system (Fig. 2B; Fig. S2B), we predicted that the cell type driving this commitment would be progenitor-like and characterized by negative covariance with reproductive cell types.

**Fig. 3.**
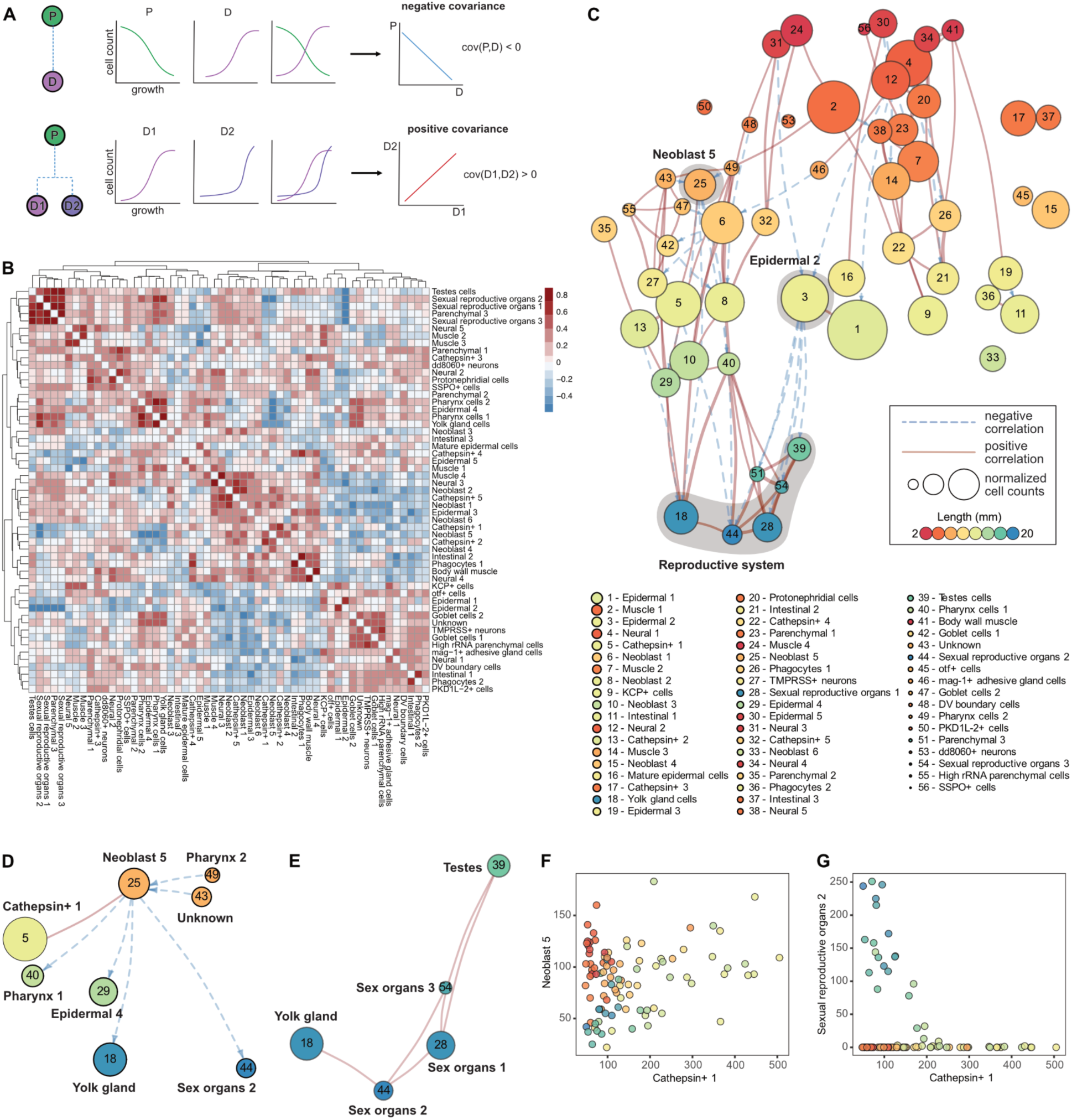
Identifying the key cell types driving the two reproductive fates. (**A**) Schematics of the covariance signatures used to infer developmental relationships. Top: a progenitor (P) and its descendant (D) show negative covariance, as the progenitor pool is depleted while the descendant accumulates during growth. Bottom: two descendants (D1, D2) of a common progenitor show positive covariance, as both accumulate in parallel. (**B**) Size-corrected cell-cell correlation matrix shown as a heatmap with hierarchical clustering; color indicates the strength and sign of the correlation. (**C**) Network representation of the size-corrected covariance structure; only significant edges are shown (threshold: mean + 1.75 s.d.). Edges denote positive (solid red) or negative (dashed blue) correlations, node size reflects normalized cell counts, and node color indicates mean body length. Numbered nodes correspond to the cell types listed below. (**D**) Subnetwork centered on Neoblast 5. (**E**) Subnetwork for the reproductive cell types. (**F**) Per-worm abundance of Neoblast 5 versus *Cathepsin*⁺ 1, with each point colored by body length. (**G**) Per-worm abundance of Sexual reproductive organs 2 versus *Cathepsin*⁺ 1, colored as in (F).

We first analyzed raw cell-cell correlations and found that >95% of the covariation could be explained by size alone, grouping most cell types into highly correlated clusters unlikely to represent genuine developmental relationships (Fig. S3A). Accounting for body size in the model removed this confounder and exposed a richer interaction structure that recapitulated known developmental relationships (Fig. 3B). Among the cell types with multiple negative associations are neoblast subpopulations (Fig. S3B), which, as the only dividing cells in planarians, are expected to give rise to all somatic and germline lineages (*13*, *33*, *34*). The strongest positive associations were found among cell types comprising the sexual apparatus, including a subpopulation of parenchymal cells (Fig. 3B). We visualized all significant cell-cell dependencies in the size-corrected model as a graph and found positive edges (coordinated growth) occurred far more frequently than negative (progenitor-like) ones across the network (see Fig. 3C, S3C). The scarcity of negative edges indicates this signature does not arise spuriously from the compositional nature of the data, but instead marks a small set of cell types with progenitor-like behavior. While progenitor-like behavior is limited to a few cell types, we reasoned that a progenitor contributing to multiple lineages should have many connections and be dominated by negative edges (*32*). Counting the total edges per cell type (Fig. S3D), we found that a single neoblast subpopulation, Neoblast 5, was one of the most connected nodes in the network, and that all but one of its edges were negative (Fig. 3D). Its negatively correlated partners include cell types of the sexual reproductive system, which are themselves positively associated with the rest of the reproductive apparatus (Fig. 3E). We confirmed this cell abundance-based link using a transcriptome-based Jensen-Shannon distance, reasoning that lineally related cells would show higher gene expression similarity (Fig. S3H). Neoblast 5 cells showed strong transcriptional similarity to *Cathepsin*^+^ 1 and yolk gland cells, but the strongest reproductive cell type suggested by the cell covariance analysis was not among its nearest transcriptional neighbors, suggesting that Neoblast 5 descendants may pass through intermediates before defining terminal reproductive organs. The negative associations of Neoblast 5 were not limited to sexual reproductive cell types. They also extended to non-reproductive populations, such as Pharynx cells 1 and Epidermal 4 (Fig. 3D), which are transcriptionally distant from Neoblast 5 (Fig. S3H). This pattern is consistent with indirect rather than direct lineage relationships. Together, these patterns are consistent with a single neoblast subpopulation contributing to *de novo* production of reproductive organs, acting upstream of sexual reproductive fate.

Given its predicted role in preceding a whole-animal reproductive fate decision, we examined how Neoblast 5 abundance varied along the growth trajectory. Neoblast 5 cells were highly enriched in small, pre-reproductive worms and declined in abundance as the reproductive system emerged in larger animals (Fig. S3F), consistent with a progenitor pool that is consumed as the sexual lineage is built. We next asked whether Neoblast 5 was also positioned with respect to the asexual fate. In contrast to its largely negative, progenitor-like connections, Neoblast 5 had a single significant positive connection, with *Cathepsin*^+^ 1 cells, a cell type highly enriched in worms at the trajectory split point (Fig. S3F-G). Because *Cathepsin*^+^ 1 cells peak before either reproductive commitment, this positive association indicates that Neoblast 5 co-occurs with the cellular state at the branch point rather than differentiating into an asexual lineage. Consistent with this, comparing their abundances revealed that Neoblast 5 cells remained steady as *Cathepsin*^+^ 1 cells peaked and then declined as the worm grew (Fig. 3F). Plotting *Cathepsin*^+^ 1 abundance against a representative sexual cell type revealed a size-driven split (Fig. 3G) reminiscent of the bifurcation in the trajectory (Fig. 2C).

Together, these analyses position Neoblast 5 as a candidate progenitor of the sexual lineage that is most abundant at the trajectory branch point, consistent with a pool poised at the decision between fates rather than committed to either.

### Temperature biases the reproductive fate of *P. morgani*

The reproductive behavior of *P. morgani* aligns with seasonal shifts in temperature: sexual populations emerge in winter, whereas asexual populations predominate throughout the remainder of the year (*10*). We wondered whether this documented seasonal behavior could be represented using the reproductive trajectory resolved from our cell composition analysis in *P. morgani* (Fig. 3).

First, to recapitulate this seasonal effect under controlled laboratory conditions, we performed a growth experiment spanning temperatures from 10°C to 22.5°C in 2.5°C increments. Worms were fed and imaged weekly to track body length, fission events, and the onset of sexual maturity (Fig. 4A). Based on established knowledge of sexual maturation in other planarian species (*24*, *35–39*), we used the appearance of a fully developed gonopore as a morphological indicator of sexual maturation. Consistent with natural seasonal patterns, worms reproduced exclusively sexually at 10°C and predominantly asexually at 22.5°C, with both reproductive modes observed at intermediate temperatures (Fig. 4B). Given that body size is a known determinant of sexual maturation, we next identified the minimum body size required for reliable gonopore detection. We found that, except at 22.5°C, worms matured only upon reaching a threshold size of ∼15 mm at all temperatures. At 22.5°C, most individuals (n = 49), with one exception, failed to reach this size by the end of the experiment (Fig. S4A-B). This result suggests *P. morgani’s* seasonal modulation of reproductive mode is primarily driven by temperature, which shapes reproductive fate, at least in part, by modulating growth.

**Fig. 4.**
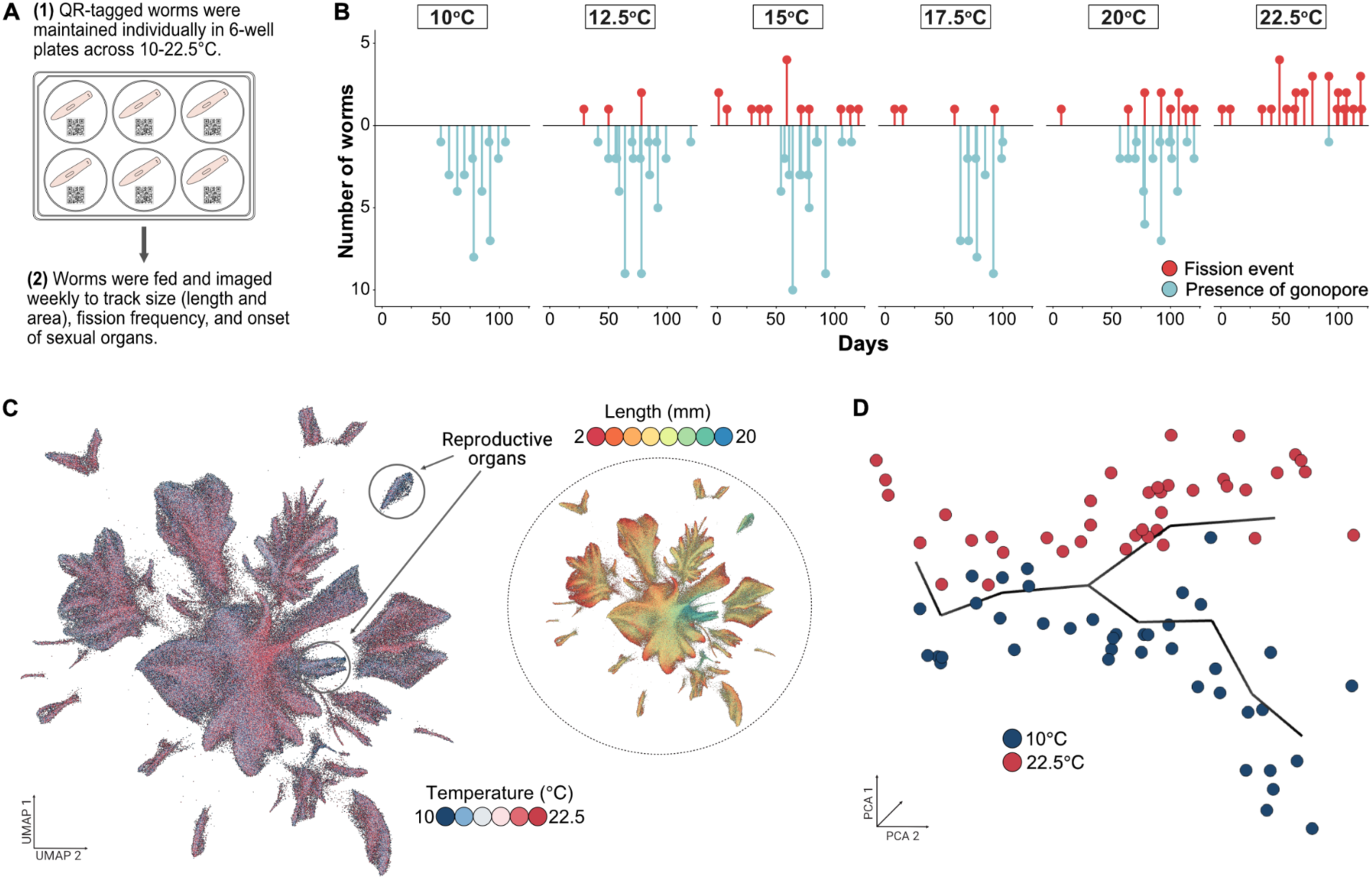
Effect of temperature on the reproductive fate of P. morgani. (**A**) Experimental workflow for assessing the effect of temperature on reproductive fate. (**B**) Tick plots showing the timing and frequency of fission events (red) and gonopore formation (blue) across the temperature gradient. Each line represents a single event; the x-axis spans the full duration of the experiment (days). Three independent replicates were performed, with 10 animals per temperature condition for the first replicate and 20 animals per temperature condition for the remaining replicates. (**C**) UMAP embedding colored by temperature treatment. Inset: the same embedding colored by body length, revealing the covariation of temperature and size across the dataset. (**D**) Growth trajectory inferred from the temperature atlas by pseudotime ordering (Monocle3) through points in PCA space, where each point represents a single worm colored by temperature treatment. Worms from the two temperature extremes (10°C and 22.5°C) segregate onto opposing branches, corresponding to exclusive sexual and asexual fates, respectively.

We next sought to exploit this temperature-dependence to directly bias the facultative reproductive trajectory. To measure this effect at high-resolution, we generated an additional single-cell atlas resolving both size and temperature effects. Worms were cultured under the same temperature regimen used in the growth experiment and sampled over a size range comparable to our initial, single-temperature atlas (Fig. S4C). Following quality control, this dataset comprised approximately 580,000 cells from 212 individuals, ranging from 2-20 mm in length across six temperature conditions (Fig. 4C; Fig. S4D). Cell-type identities were assigned via automated label transfer using the original size atlas as a reference (Fig. S4E-F). By applying the same trajectory analysis as previously described (Fig. 2C), we recapitulated the bifurcation observed in the size atlas, with the developmental trajectory again splitting into two branches, with more worms occupying the asexual branch than observed previously at 17.5°C (Fig. 4D; Fig. S4G). This stronger divergence was driven predominantly by the two temperature extremes (10°C and 22.5°C), which strictly correspond to sexual and asexual fates (Fig. 4B, D). Consistent with phenotyping results, intermediate temperatures (12.5°C to 20°C) produce worms occupying both branches, with the number of worms on the asexual branch increasing with temperature (Fig. S4G-H). Temperature extremes, therefore, engage the identical bifurcating developmental program that governs the size-dependent reproductive decision at the intermediate 17.5°C, indicating a shared cellular mechanism.

### A cold-activated progenitor couples temperature to reproductive fate

We established Neoblast 5 as a primary driver of reproductive fate during growth (Fig. 3D), but we wondered whether this same progenitor population could mediate the temperature-dependent reproductive plasticity of *P. morgani*. Our findings yield a specific prediction: if Neoblast 5 cells differentiate into sexual cell types only at permissive cooler temperatures, this differentiation should be suppressed in warmer environments, leading to accumulation of this progenitor pool. To test this hypothesis, we examined cell types whose abundance was associated with high temperatures and found that Neoblast 5 showed the strongest positive correlation (Fig. S5A). We then performed differential cell abundance analysis across the temperature gradient, identifying 38 cell types that showed significant temperature effects after accounting for size (Fig. S5B). Consistent with our prediction, Neoblast 5 was among the cell populations that increased most substantially with temperature, showing peak abundance at 22.5°C (Fig. 5A). We confirmed that the progenitor’s temperature-dependence persists even after accounting for body size (Fig. S5B), ruling out the possibility that temperature modulates reproductive solely via effects on growth. Accumulation of Neoblast 5 was mirrored in the depletion of its sexual descendants: the ratio of reproductive cell types to progenitors within individual worms fell sharply as temperature increased (Fig. 5B-C). A progenitor pool that accumulates while its descendant lineages vanish suggests suppression, rather than intrinsic reduction, of differentiation at warmer temperatures.

**Fig. 5.**
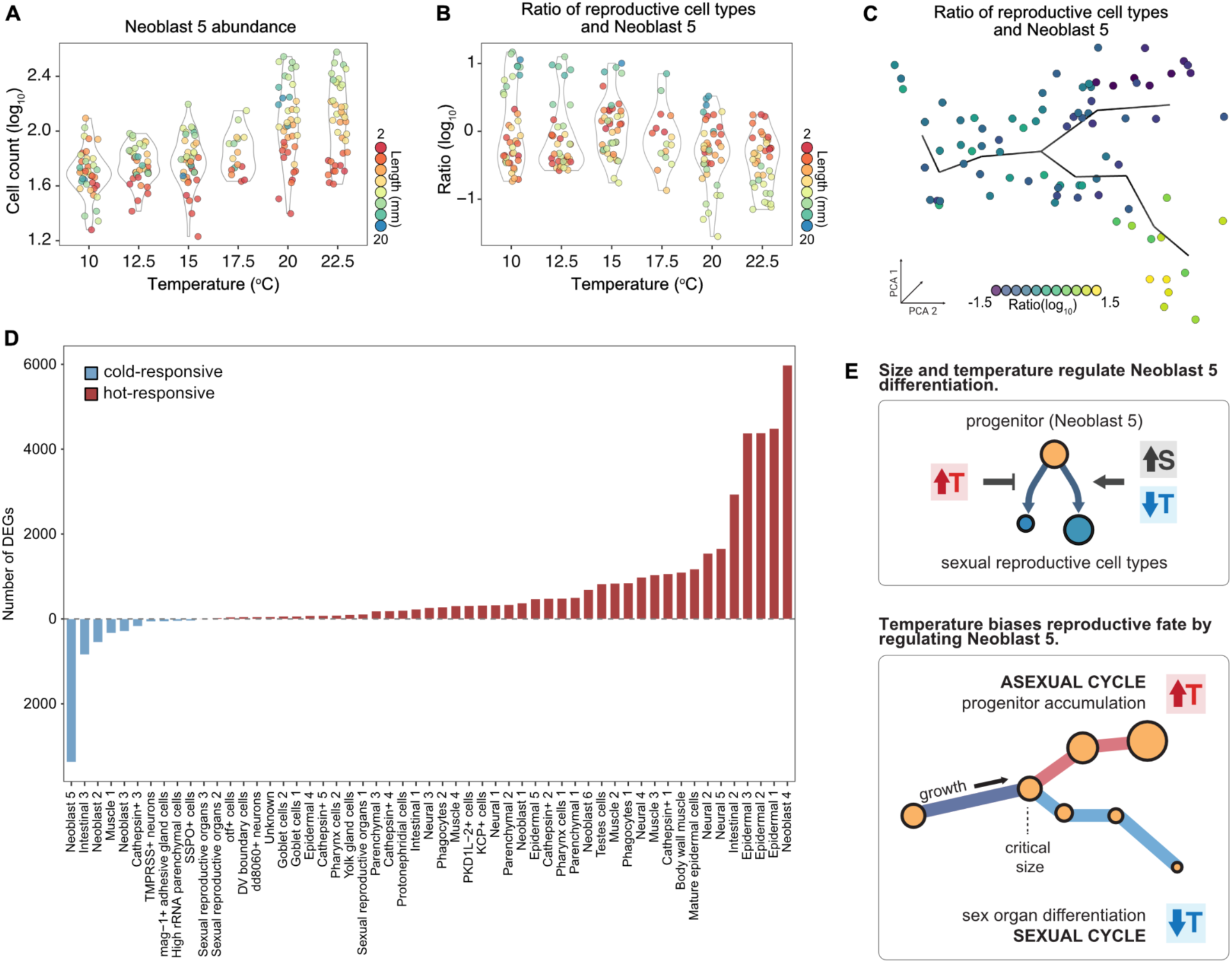
A cold-sensing neoblast population couples temperature to reproductive fate across the natural temperature range of P. morgani. (**A**) Neoblast 5 abundance (log₁₀ relative cell count) across the temperature gradient (10-22.5°C). Each point represents a single worm and is colored by body length. (**B**) The ratio of reproductive system cell counts to Neoblast 5 abundance (log₁₀) across the temperature gradient, with points colored by body length as in (A). (**C**) The same ratio plotted along the growth trajectory inferred for the temperature atlas (PCA space; Monocle3), with each worm colored by its ratio (log₁₀). (**D**) Temperature-dependent changes in the number of differentially expressed genes (DEGs) per cell type, ranked and colored by response direction (cold-responsive: blue; warm-responsive: red). (E) Working model for Neoblast 5 as a cold-activated progenitor that couples body size and temperature to reproductive fate. Top: body size and temperature act on Neoblast 5 differentiation. Larger size (↑S) and lower temperature (↓T) transcriptionally activate the pool and promote its differentiation into sexual reproductive cell types, whereas higher temperature (↑T) keeps the pool transcriptionally inactive and suppresses its differentiation. Bottom: the same logic placed along the growth trajectory. Beyond a critical body size the trajectory splits; at high temperature the progenitor pool remains transcriptionally inactive and accumulates, biasing worms toward the asexual branch, whereas at low temperature the pool becomes transcriptionally active and is consumed as it differentiates to construct the reproductive system, biasing worms toward the sexual branch. Circle size reflects relative Neoblast 5 abundance.

To uncover the molecular basis of this suppression, we took advantage of underlying gene expression data and asked whether temperature-dependent expression differences reflected effects we identified at the level of cell abundance. Specifically, we compared cold vs. warm transcriptomes within each cell type to define a temperature-response that is independent from growth or cell identity programs. From this unbiased analysis across all cells, Neoblast 5 emerged as one of a small subset of cell types exhibiting a large number of cold-responsive genes and was the only cell type among these linked to the reproductive system (Fig. 5D). Gene Ontology (GO) analysis of cold-responsive genes in Neoblast 5 revealed an enrichment of DNA-binding-related terms. This finding is consistent with the activation of a differentiation-associated transcriptional program (Fig. S5E). Thus, Neoblast 5 abundance and transcriptional activity move in opposite directions in response to temperature. At 10°C, the progenitor pool becomes transcriptionally active and is consumed as it differentiates into reproductive lineages. At 22.5°C, the progenitor pool remains transcriptionally inactive and accumulates due to the blockage of its differentiation into sexual lineages. This decoupling of abundance from activity reconciles the high Neoblast 5 abundance at asexual-promoting temperatures with its definitive role in constructing the sexual reproductive system (Fig. 5D-E). Collectively, these results identify a single, unifying cellular mechanism centered on Neoblast 5 that dictates both size-driven and temperature-induced reproductive decisions. By gating the differentiation of a cold-activated progenitor, temperature directly couples environmental conditions to reproductive fate.

## Discussion

Despite the cellular and molecular complexity underlying the *P. morgani*’s life-cycle, we find that its reproductive fate is controlled by a single, cold-activated progenitor population. Neoblast 5 cells couple body size and temperature to a binary developmental outcome via differentiation status (Fig. 5E). Under cold, sexually-promoting conditions, Neoblast 5 cells are transcriptionally activated and differentiate into reproductive lineages that form the sexual apparatus *de novo*. Under warm, asexually-promoting conditions, progenitors accumulate while their sexual descendants are depleted, a signature of suppressed differentiation rather than intrinsic progenitor loss. This architecture provides a parsimonious explanation for how one animal, from the same genome and the same stem cell reservoir, produces two distinct reproductive phenotypes.

High throughput profiling of cellular complexity by individually resolved sci-RNA-seq allowed us to reconstruct a whole-animal growth trajectory for a new model system in a single experiment. Building this trajectory directly from data over hundreds of individuals revealed a fate decision with defined geometry: a bifurcation at intermediate body size that separates worms into sexual and asexual branches before any anatomical difference is apparent. The sexual branch is marked by the progressive appearance of testes, yolk gland cells, and other reproductive cell types, while the asexual branch is characterized by enrichment of subpopulations of parenchymal, epidermal, and pharynx cells together with the absence of reproductive lineages. That the same bifurcation in cell composition is recapitulated by temperature treatments, with cold and warm temperatures segregating worms to opposite branches, confirms that size and temperature act on the same underlying cellular program rather than independent mechanisms.

Systematic analysis of cell-cell growth interactions positions Neoblast 5 as a candidate regulator, operating at the intersection of alternative reproductive fates. Among cell types with proliferative and differentiation potential, it is the most connected node in the size-corrected cell-cell interaction network, with almost exclusively negative edges, the expected signature of a progenitor whose abundance falls as its descendants accumulate (*32*, *40*). Surprisingly, we also identified Epidermal 2 cells as showing progenitor-like behavior for sex organs (Fig. 3C). This mechanism must be indirect given these cells are post-mitotic. The critical position of Neoblast 5 cells is reflected in their capacity for temperature-sensing; these cells exhibit one of the most pronounced cold-responsive transcriptional programs in the worm and are the only cold-responsive cell type directly linked to the reproductive system. The sensitivity of Neoblast 5 cells over the full natural temperature range of *P. morgani* argues against a generalized stress response and in favor of a regulatory mechanism that acts specifically to control reproductive differentiation. In a model system with no established lineage tracing or perturbation tools, predicting developmental relationships among cells using scalable single-cell profiling and statistical analysis is a powerful approach, but direct validation of the inferred descendant relationships is an important next step. Further, the molecular processes licensing Neoblast 5 activation at low temperatures, whether through neuroendocrine, metabolic, or niche-mediated signals, likewise remains an open question, and the cold-responsive transcriptional signature identified here nominates candidate regulators for future investigation.

These findings would not have been accessible without individual-level resolution. The bifurcating trajectory is a property of the distribution of individual worms in compositional space, and pooling cells across animals would have collapsed it into a population average. More critically, the decoupling of Neoblast 5 abundance from its transcriptional activity, high and quiescent at warm temperatures and low and active at cold, can only be detected when compositional and transcriptional states are measured in the same individual. This decoupling is the most counterintuitive feature of the mechanism, with implications beyond this system: a progenitor pool whose size is inversely related to the lineage it produces would be misread as inactive by any analysis that treats abundance as a proxy for productive output. We term this phenomenon differentiation gating, a configuration in which a progenitor is held transcriptionally inactive under non-permissive conditions and released into differentiation only when a specific signal is present.

Differentiation gating is a recurring feature of regenerative systems. The closest parallel is the hair follicle, where bulge stem cells persist as an abundant, quiescent pool during the resting phase and are activated into proliferation and differentiation at the onset of the growth phase, so that progenitor abundance is highest precisely when productive output is lowest (*41*, *42*). This cycle can be regulated by external environmental signals, such as temperature and light. These signals are relayed to the stem cell niche through the sympathetic nervous system (*43*, *44*). These examples offer a precedent for how an environmental cue might couple to a quiescent progenitor pool through a systemic relay rather than a cell-autonomous sensor, and nominating candidate mechanisms for the cold-licensed activation of Neoblast 5. In *P. morgani*, the effect of differentiation gating is harnessed for facultative reproduction, selecting between two alternative fates of the body rather than driving a cyclical regenerative program. We expect that this cellular mechanism operates in other species, but identifying such architectures will require measuring both cell abundance and transcriptional state across individuals. New approaches for scalable, individually resolved single-cell profiling brings life-cycle trajectory inference within reach, and may uncover new mechanisms of phenotypic plasticity for unconventional species and natural contexts alike.

## Acknowledgments

We thank members of the Vu and Dorrity groups for their input and discussions. We thank the Genomics Core Facility and Advanced Light Microscopy Facility at EMBL Heidelberg for support with sequencing and imaging. We thank Dr. Amrutha Palavalli for her advice during the optimization of FISH for *P. morgani* and her generous help with nuclei extractions and imaging. We thank Eric Bormann for his help with optimizing the in situ protocol, respectively. We thank Daniel Stoga for maintaining the planarian stocks. We thank Dr. James Cleland for collecting the original colony of *P. morgani*. We thank Dr. Alejandro Sánchez Alvarado for his support leading up to and throughout this work. We thank Dr. Aissam Ikmi and Dr. Jordi van Gestel for their feedback on the manuscript. We declare the use of AI tools, including ChatGPT and Claude, to assist with debugging and refining code for some analyses, and to check grammar and improve clarity of the text. AI tools were not used to generate the original content of the manuscript.

## Author contributions

Conceptualization: GV, HT-KV, MWD

Methodology: GV, AMMF, HT-KV

Resources: GV, KM, AUMK, CT, ER, SR

Investigation: GV, EL, AAMF, DY

Formal analysis: GV, AG

Visualization: GV

Supervision: HT-KV, MWD

Writing – original draft: GV, HT-KV, MWD

Writing – review & editing: GV, HT-KV, MWD with input from all co-authors

## Diversity, equity, ethics, and inclusion

The authors declare that no aspects of the Diversity, equity, ethics, and inclusion criteria were applicable to this study.

## Data, code, and materials availability

The raw sequencing data generated and the processed FASTQ files used in this study have been deposited in the European Nucleotide Archive (ENA) at EMBL-EBI (accession number pending). Raw image datasets and the corresponding metadata are deposited in the BioImage Archive at EMBL-EBI under (accession number pending). Processed single-nucleus datasets and a user-friendly data browser will be made available at our website: <pending url>. All original code, including notebooks for performing data processing, statistical analysis and generating plots, has been deposited at Github and will be made publicly available as of the date of publication: <pending url>.

## Supplementary Materials for

## Materials and Methods

### Worm maintenance and sampling

The original colony of *P. morgani* was kindly shared with us by the lab of Dr. Jochen Rink. The worms were collected in Point of Rocks, Virginia (39.273359, −77.536762). For the size atlas, worms of varied body sizes were acclimated and maintained at 17.5°C in Montjuïc planarian water (*45*). For the size-temperature atlas, worms of comparable body size (∼2 mm) were acclimated to the indicated temperatures and maintained in Montjuïc planarian water. Worms were grown at their respective acclimation temperatures for ∼6 months, during which samples for nuclei preparation were collected weekly. Worms were fed homogenised calf liver once per week, and those selected for sampling were starved for more than 1 week before being snap-frozen in liquid nitrogen and stored at −70°C.

### Imaging and image analysis pipeline for planarian body size measurement

For the single-cell atlases, worms selected for sampling were imaged prior to snap-freezing in order to measure their size.To ensure accurate body length measurements, MP4 videos were recorded for each worm to capture individuals in their maximally stretched state. These videos were subsequently analysed using a custom-developed image analysis pipeline (Fig. S1A) (https://git.embl.org/grp-cba/planarian-body-size-measurement). Raw videos were first processed into a series of JPEG images subsampled at a rate of 2 frames per second (fps). YOLOv8 (You Only Look Once Version 8, (https://github.com/ultralytics/ultralytics), a state-of-the-art deep learning-based object detection model, was trained to detect two classes: worm and a QR code. The training dataset was based on images of *Phacogata morgani* and *Schmidtea mediterranea* of various body sizes. Each image also included a unique QR code associated with the worm. The worms and the QR codes in the training images were labeled with rectangular bounding boxes using YoloLabel (https://github.com/developer0hye/Yolo_Label). The QR codes were used for tracking individual worms over multiple imaging sessions. Since such tracking was not required for worms sampled for the single-cell atlases, the trained object detection model was instead used solely to identify individual worms in each image and generate a bounding box around the detected animal. The region of interest (ROI) defined by each bounding box for worms was then cropped from the original image and passed to SAM (Segment Anything Model) for segmentation (https://github.com/facebookresearch/segment-anything). From this segmentation, morphological features including the feret’s diameter (measurement for body size), area, circularity and solidity were then extracted using the *regionprops* function of the scikit-image Python library. The frames were filtered on the basis of the worm’s circularity and solidity to remove erroneous segmentation. The frame containing the worm with the largest area and length-to-width ratio was used for the size measurement.

For tracking individual worms over multiple imaging sessions as required for the growth experiment, a unique QR code was associated with each worm stuck to the bottom of the well and imaged with the worm.For these experiments, the same trained model was used to detect and fit independent bounding boxes around both the worm and its associated QR code within each image. The QR code information was then read out using an R script and the decoded information was added to the corresponding frame’s CSV file.

### Growth experiment

Worms of approximately equal body size (6-8 mm) were acclimated to different temperatures used in the growth experiment and maintained individually in wells of 6-well plates. At a later stage of the experiment due to increasing body size, worms were transferred to individual 9 mm petri plates. Each well and, therefore, worm was assigned a unique identifier in the form of a QR code affixed to the bottom of the well, which was captured during video recording for body size measurement. Worms were fed homogenized calf liver and imaged weekly to monitor changes in body size. Body size measurements were performed as described previously. Worms that underwent fission events or died during the experiment were excluded from size analysis, and fission fragments were removed from the wells. Gonopore formation and fissioning events were checked weekly during the experiment and verified by re-examining the recorded videos after completion of the experiment.

### RNA isolation and sequencing for de novo transcriptome assembly for *P. morgani*

A clonal population of worms was used to extract RNA for the transcriptome. We sampled both regenerating and intact worms, which were starved for more than a week prior to RNA extraction. Sampled worms were divided and snap-frozen in 1.5 mL DNA LoBind tubes (Eppendorf, EP0030108051). Frozen worms were homogenized in a mixture of 1 M dithiothreitol (DTT) (Biomol, D8070.10) and TRIzol (Thermo Fisher, 15596026) reagent at a 1:9 ratio (100 μL of 1 M DTT and 900 μL of TRIzol reagent), using 5-6 zirconia beads (Carl Roth, N039.1) in a Bead Ruptor Elite (2 cycles; 7.45 m/s for 45 seconds, with a dwell time of 45 seconds). The solution was incubated at room temperature and transferred to 2 mL Phasemaker tubes (Thermo Fisher, A33248). Chloroform (0.2 mL) was added to the homogenate, and the tube was shaken vigorously for 15 seconds. The tube was incubated at room temperature for 3 minutes and spun at 12,000 × g for 15 minutes at 4°C. The aqueous phase at the top was removed and transferred to a fresh tube. The aqueous phase was mixed with 0.25 mL of RNase-free isopropanol (Thermo Fisher, 327272500), followed by 0.25 mL of a high-salt precipitation solution (0.8 M sodium citrate, 1.2 M NaCl) per 1 mL of TRIzol reagent added, and the solution was vortexed for a few seconds until the precipitate disappeared. The mixture was column-purified and eluted in 30 μL of DNase/RNase-free water. RNA extracted from multiple samples was pooled, and concentration and RNA integrity were determined using a NanoDrop and a TapeStation 4200 (Agilent). Libraries for short-read Illumina and long-read PacBio Iso-Seq sequencing were prepared and sequenced at the Genomics Core Facility at EMBL Heidelberg.

### *de novo* assembly of *P. morgani* transcriptome

*P. morgani* transcriptome was assembled by combining long-read and short-read sequencing strategies. Short-read RNA-seq data were assembled de novo using Trinity (*46*, *47*) (v2.15.1) with strand-specific (RF) paired-end library settings. Long-read PacBio HiFi data were processed using the IsoSeq3 (v3.8.1) workflow: reads were trimmed with ‘lima’, artifacts were removed and poly-A tails required using ‘isoseq refine’, and high-quality isoforms were clustered with ‘isoseq cluster’. Within-assembly redundancy in the IsoSeq transcripts was reduced using CD-HIT-EST (*48*, *49*) at 95% nucleotide identity. The two assemblies were concatenated and further collapsed with CD-HIT-EST at 95% global identity to produce a non-redundant combined assembly. RNA-seq reads were then grouped into genes/clusters using Salmon (*50*) and Grouper (*51*). Sequences were renamed to reflect cluster membership (prefix pmor.kc3, gene cluster identifier gcXXXXXX). Transcripts identified as foreign contamination by NCBI FCS-GX (v0.4.0) (*52*) were removed, and transcripts shorter than 300 bp were discarded.

For scRNA-seq analysis, a non-redundant set of transcripts was generated by taking the most representative isoform/transcript from each cluster as determined by OMA (*53*, *54*).

### cDNA synthesis

Total RNA was isolated from large sexual P.morgani worms using the protocol described above. First-strand cDNA synthesis was performed in 1.5 mL DNA LoBind tubes (Eppendorf, EP0030108051). 1 μL of dT_20_ oligo, 6 μL of total RNA and 1 μL 10 mM dNTP mix were combined and brought to a final volume of 13 μL using nuclease-free water. This mixture was heated to 65°C for minutes in a thermomixer and then immediately chilled on ice for 1 minute. Following brief centrifugation, 4 μL 5X First-Strand Buffer, 1 μl of SuperScript™ III RT (200 units/μl) (Thermo Fisher, 18080093), 1 μl 0.1 M DTT and 1 μl Recombinant RNase Inhibitor were added to the reaction. The contents were mixed by pipetting up and down gently. The tube was incubated at 50°C for 45 minutes followed by incubation at 70°C for 15 minutes. Finally to remove RNA, 0.4 μL (2 units) of RNase H (NEB, M0297L) was added to the tube and incubated at 37°C for 20 minutes. cDNA concentration and quality was checked using a nandrop. The cDNA sample was stored at −70°C.

### Candidate gene selection for *in situ* hybridization

Cell-type-specific marker genes were identified using the *top_markers()* function in Monocle3. For each of the 55 cell types, the top 25 markers were computed and ranked by their specificity score. Candidate markers were selected from the ten highest-ranking genes for each cell type and complemented with established planarian cell-type markers from previously published studies. Transcript sequences for the selected candidate markers were retrieved from the transcriptome.

### Riboprobe synthesis

Primers were designed against target transcript sequences using a combination of manual design and Primer3plus (https://www.primer3plus.com/) while maintaining consistent design parameters across all targets (Supplementary Table S5). Riboprobe synthesis was carried out in three sequential steps involving two rounds of PCR followed by an vitro transcription reaction. In the first step target sequences were amplified by touchdown PCR while simultaneously incorporating primer overhangs required for subsequent template generation. The resulting amplicons were then used as templates for the second PCR reaction, during which a T7 RNA polymerase promoter sequence was added. These T7-tagged PCR products served as templates for the final in vitro transcription step. The touch-down PCR was performed using the NEB *Taq* polymerase (NEB, M0273) protocol for 50 μL reactions (https://www.neb.com/en/protocols/taq-dna-polymerase). The following PCR cycle conditions were used for the touch-down PCR: 95°C, 30 seconds; 10 cycles of 95°C, 30 seconds, 63°C - 53°C for 1 minute (1°C decrease per cycle) and 68°C for 1 minute 30 seconds; 25 cycles of 95°C, 30 seconds, 53°C for 1 minute and 68°C for 1 minute 30 seconds; 68°C for 5 minutes; 10°C hold. The second PCR for DNA template synthesis followed the same protocol as the first for the PCR reaction mix. The following PCR cycle conditions were used for the second PCR: 94°C for 5 minutes; 30 cycles of 94°C for 30 seconds, 54°C for 15 seconds and 68°C for 3 minutes; 68°C for 10 minutes; 10°C hold. PCR products from both amplification steps were purified using a PCR purification kit (Qiagen, 28106). Purified DNA templates were used for in vitro transcription. 500-700 ng of the purified DNA template was mixed with 2 μL of 10x Trans buffer (0.1M MgCl_2_, 0.4M Tris pH 8.0, 0.1M DTT, 20mM spermidine (Sigma, S2626-1G), RNAse-free water), 10x RNA Labeling mix DIG (Roche, 11277073910), 0.5 μL RNAse inhibitor (ThermoFisher, EO0382), 2 μL T7 polymerase (PepCore, EMBL Heidelberg), 0.3 μL inorganic pyrophosphatase (ThermoFisher, EF0221) and brought up to 20 μL using nuclease-free water. Transcription reactions were incubated overnight at 37°C. To remove the DNA template, 1 μL DNase I (NEB, M0303S) was added and the reaction was incubated for an additional 45 min at 37°C. The resulting RNA was precipitated by adding 0.5 volumes of 7.5M ammonium acetate, followed by 2.5 volumes of ice-cold 100% ethanol. Samples were vortexed thoroughly and incubated at −70°C for at least 30 minutes. Following precipitation, samples were centrifuged at maximum speed for 30 minutes at 4°C and the supernatant was carefully removed. The resulting pellet was washed twice with 1 mL of ice-cold 75% ethanol. For each wash samples were vortexed, centrifuged at maximum speed for 5 min at 4°C, and the supernatant was discarded. After the final wash, the pellet was briefly air-dried and resuspended in 100 μL of Formamide (Carl Roth, P040.2). Riboprobes were stored at −70°C until further use.

### Sample fixation for *in situ* hybridization

Worms were incubated in ice-cold 1× Montjuïc water at 4°C for 30 minutes in 6-well plates kept on ice. Worms were then incubated in a nitric acid relaxant solution (0.2 M MgSO_₄_, ∼1.36% (v/v) HNO_₃_, H_₂_O) for 5 minutes, before the solution was replaced with a mucus-removal solution (0.5% (w/v) N-acetyl-L-cysteine (Carl Roth, 4126.2), pH adjusted to 7.25 using 1 M NaOH). Worms were incubated in the mucus-removal solution for 10 minutes before it was replaced with PFA fixative (4% PFA in PBSTx). Worms were incubated in the fixative for 1 hour, followed by 2 washes with PBSTx for 5 minutes each. Fixed worms were dehydrated in 50% methanol for 5 minutes and then transferred to 100% methanol. Fixed worms were stored in 100% methanol at −20°C.

### Whole-mount *in situ* hybridization

Fixed worms were transferred to a 6-well plate, and 100% methanol was replaced with 2 mL of 50% methanol for 5 minutes. Thereafter, the worms were washed with 2 mL of 1× SSC containing 0.1% Tween-20 per well for 5 minutes. Worms were then bleached for 2 hours using freshly prepared bleaching solution (0.5× SSC, 1.2% H_₂_O_₂_, 5% formamide, H_₂_O). Post-bleaching, worms were again washed with 1× SSC containing 0.1% Tween-20 (2 mL/well) for 5 minutes, followed by a wash with PBSTx (30 mL of 10% Triton X-100 in 100 mL of 10× PBS, diluted to 1 L with H₂O) (2 mL/well) for 5 minutes. Subsequently, PBSTx was replaced with proteinase K solution (0.1% SDS, 2 μg/mL proteinase K, 1× PBS). Worms were incubated in proteinase K solution for 30 minutes at room temperature, then fixed again with the PFA fixative (4% PFA, PBSTx) for 5 minutes.

Fixation was followed by multiple washing steps: first with PBSTx for 5 minutes, then with a 1:1 mixture of PreHybe (50% formamide, 5× SSC, 1× Denhardt’s, 100 μg/μL heparin, 1% Tween-20, 1 mg/mL torula yeast RNA, 50 mM DTT) and PBSTx for 10 minutes (2 mL/well). Worms were then incubated in preheated PreHybe solution (2 mL/well) at 58°C in a Boekel Scientific Shake ‘N Bake Rocking Hybridization Oven for 1 hour (PreHybe was preheated to 58°C in a water bath). In parallel, riboprobes were diluted 1:3000 in 3 mL of Hybe solution (50% formamide, 5× SSC, 1× Denhardt’s, 100 μg/μL heparin, 1% Tween-20, 0.25 mg/mL torula yeast RNA, 50 mM DTT, dextran sulfate) and kept at 58°C in the water bath for 1 hour. To conclude the first day of the protocol, PreHybe was replaced with the preheated Hybe-riboprobe mix (1 mL/well), in which the worms were incubated overnight.

The following day, worms underwent multiple washes: first with preheated WashHybe (50% formamide, 0.5% Tween-20, 5× SSC, 1× Denhardt’s) at 58°C for 1 hour (4 mL/well), followed by a wash with a combination of WashHybe and the 1st wash solution (2× SSC, 0.1% Tween-20) mixed in a 1:1 ratio (4 mL/well) at 58°C for 1 hour. This was followed by a wash with only the 1st wash solution at 58°C for 1 hour (4 mL/well), then a wash with the 2nd wash solution (0.2× SSC, 0.1% Tween-20) at 58°C for 1 hour (4 mL/well). Plates were cooled to room temperature before worms were washed with PBSTx for 15 minutes at room temperature (4 mL/well). Following all the washes, the worms were incubated in a blocking solution (2 mL/well) (5% horse serum, 1% Roche Western Blocking Reagent, PBSTx). Worms were incubated in the antibody solution (1 mL/well) (αDIG-POD 1:2000 in blocking solution; filtered using a Millex PVDF 0.22 μm membrane syringe filter) overnight at 4°C.

The following morning, worms were washed with PBSTx for a total of 2 hours, changing the solution 3 times during this period. This was followed by 30 minutes of incubation with the tyramide solution (4-IBPA at a 1:1000 dilution, Cy3 at a 1:1000 dilution, 30% H_₂_O_₂_ at a 1:5000 dilution, all in TSA buffer (2 M NaCl, 0.1 M boric acid, pH 8.5), filtered using a Millex PES 0.22 μm membrane syringe filter). The tyramide solution was then replaced with PBSTx for 10 minutes at room temperature (4 mL/well). Samples were subsequently stained with DAPI overnight (1:5000 DAPI in PBSTx) at 4°C.

This protocol was followed by an additional treatment to reduce autofluorescence (generously shared by the lab of Dr. Jochen Rink). This included another wash with PBSTx for 5 minutes (2 mL/well), followed by 10 minutes of fixation with PFA. Post-fixation, worms were again washed with PBSTx for 5 minutes. To improve the signal-to-noise ratio, worms were incubated in a permeabilising detergent solution (5 g sodium deoxycholate, 50 mL 1 M Tris pH 7.5, 2 mL 0.5 M EDTA, 30 mL 5 M NaCl, 100 mL 10% SDS, made up to 1 L with dH_₂_O) for 30 minutes (2 mL/well). This was followed by one final wash with PBSTx before placing the worms in ScaleS4 (*55*) (10% glycerol, 15% DMSO, 40% sorbitol, 4 M urea, 0.1% Triton X-100) overnight. Worms were finally mounted on microscope coverslips in ScaleS4.

ISH-stained samples were imaged on an Olympus IXplore SpinSR confocal microscope, except pc2 (Fig. S1E, left), which was imaged on a Leica SP8 confocal microscope. Fiji/ImageJ (https://github.com/imagej/ImageJ.git) was used to process all the images. All images shown are maximum-intensity projections of representative results.The color of the images was set to grey before inversion.

### Preparation of nuclei suspensions from individual worms

Frozen worms were dissociated into single-nuclei suspensions following the previously published sci-RNA-seq3 protocol (*21*), with modifications to adapt it for planarians. Hypotonic lysis buffer A (1× hypotonic PBS, 3 mM MgCl_₂_, 0.3 M sucrose) was added to each tube depending on the worm size (2-3 mm: 500 μL; 5-9 mm: 1 mL; 10 mm and above: 1.5 mL), together with 8-10 zirconia beads (Carl Roth, N039.1). Samples were homogenised using a Bead Ruptor Elite (2 cycles; 2.1 m/s for 15 seconds, with a dwell time of 15 seconds). Larger animals (>12 mm) were subjected to an additional round of homogenisation. The resulting homogenate was added to an appropriate DNA LoBind tube (1.5-50 mL) (Eppendorf, EP0030108051; EP0030108078; EP0030108310; EP0030122585; EP0030122615) containing hypotonic lysis buffer A supplemented with 0.01% Igepal. The volume of lysis buffer used depended on the worm size (2-3 mm: 500 μL; 5-9 mm: 5 mL; 10-12 mm: 10 mL; 13-14 mm: 20 mL; >15 mm: 30 mL). Samples were incubated on ice for 3 minutes with occasional inversion before being filtered through a 50 μm cell strainer into fresh DNA LoBind tubes. Filtered lysates were centrifuged at 700 × g for 5 minutes at 4°C. The supernatant was carefully poured out, and the pellet was resuspended in 1 mL of 0.3 M SPBSTM + 1% DEPC (Sigma, D5758-50ML) by gentle pipetting (>10 times). Additional 0.3 M SPBSTM + 1% DEPC was added to the tubes for washing depending on the worm size (2-3 mm: none; 4-5 mm: 1 mL; 6-8 mm: 5 mL; 9-12 mm: 10 mL; >13 mm: 20 mL). Samples were centrifuged at 700 × g for 5 minutes at 4°C; the supernatant was poured out gently, and the nuclei pellet was resuspended in 1 mL of 0.3 M SPBSTM + 1% DEPC.

Nuclei for each sample were quantified by mixing 9 μL of nuclei with 1 μL of 10× dsGreen DNA stain (Lumiprobe, 20010) and counting using C-Chip counting chambers (Carl Roth, PK36.1). For worms smaller than 10 mm, the entire nuclei preparation was fixed. For larger worms, aliquots containing the desired number of nuclei were transferred to fresh DNA LoBind tubes. The volume was adjusted to 1 mL with 0.3 M SPBSTM + 1% DEPC, followed by centrifugation at 700 × g for 5 minutes at 4°C. Finally, nuclei were resuspended in 100-200 μL of 0.3 M SPBSTM + 1% DEPC for fixation. The nuclei were fixed using bis(sulfosuccinimidyl)suberate (BS3) (Thermo Fisher, 21586), an amine-reactive cross-linker dissolved in methanol at a final concentration of 1.25 mg/mL (500 μL fixative for 100 μL nuclei). The fixative was added slowly and gently mixed by pipetting. Samples were then incubated on ice for 15 minutes with occasional swirling to mix. Following fixation, nuclei were rehydrated by adding 2 volumes of 0.3 M SPBSTM dropwise while gently shaking the tube to ensure proper mixing. Samples were centrifuged at 700 × g for 10 minutes at 4°C, and the supernatant was carefully removed. The nuclei pellet was resuspended in 200 μL of 0.3 M SPBSTM by gentle pipetting (>20 times), after which the volume was brought up to 1 mL with additional buffer. Samples were centrifuged again under the same conditions, and the nuclei were resuspended in 1 mL of fresh 0.3 M SPBSTM.

Nuclei concentrations were determined by mixing 2 μL of nuclei suspension with 7 μL of 0.3 M SPBSTM and 1 μL of 10× dsGreen DNA stain. Nuclei were counted using C-Chip counting chambers (Carl Roth, PK36.1). Based on the counts obtained, nuclei were diluted in 0.3 M SPBSTM to the desired concentration and aliquoted into 8-well strip tubes. Approximately 30,000 nuclei were collected per sample for the size atlas and approximately 60,000 nuclei per sample for the size-and-temperature atlas. Aliquoted nuclei were pelleted by centrifugation at 700 × g for 10 minutes at 4°C; the supernatant was removed, and the strip tubes were snap-frozen in liquid nitrogen before storage at −70°C until further processing.

### sci-RNA-seq3 library construction

The fixed nuclei were processed using the previously published sci-RNA-seq3 protocol (*21*), with several modifications to improve reverse transcription efficiency and reduce the proportion of uncorrectable barcodes. Nuclei were thawed on ice, and approximately 20,000 nuclei per sample were distributed into individual wells across three 96-well LoBind plates (Eppendorf, EP0030129512), yielding three technical replicates per sample. A mixture of 10x dsDNAse buffer (Thermo Fisher, EN0771), dsDNase (Thermo Fisher, EN0771) and dNTPs (NEB, N0447L) was added to the nuclei in each well and incubated for 2 minutes at 37°C. This was followed by adding 2 μL of 10 mM 3-level RT primers (IDT) carrying the first index. The plates were incubated at 55°C for 5 minutes in a thermocycler (lid temperature 65°C) to inactivate the dsDNase. Maxima H Minus reverse transcriptase (Thermo Fisher, EP0753), Ribolock RNase inhibitor(Thermo Fisher, EO0382) together with 25% PEG to increase RT efficiency, was then added to each well, and the plates were incubated at 4°C, 10°C, 20°C, 25°C, 30°C, 40°C, 50°C, and 55°C, with 2-minute cycles at each temperature except 25°C (10 minutes) and 55°C (30 minutes).

Cold 0.3 M SPBSTM was added to each well, and the nuclei were gently mixed by pipetting, after which wells from all three plates were pooled. The pooled sample was centrifuged at 700 × g for 5 minutes at 4°C, and the resulting pellet was washed with 0.3 M SPBSTM. The suspension was centrifuged again under the same conditions and resuspended in 0.3 M SPBSTM. The pooled nuclei were then sonicated at low intensity for 6 seconds at 4°C using a Diagenode Bioruptor Plus.

Next, 11 μL of nuclei were distributed per well into four fresh 96-well LoBind plates, followed by 2 μL of 10 mM 3-level ligation primers (IDT) carrying the second index. T4 DNA ligase (NEB, M0202L; 2 μL) in T4 ligation buffer was added to each well, and the plates were incubated for 30 minutes at 20°C. Cold 0.3 M SPBSTM (10 μL) was then added to each well, and all wells from all plates were pooled. The pooled sample was centrifuged and washed twice with 0.3 M SPBSTM as described above to remove the ligation buffer. The nuclei were resuspended in 0.3 M SPBSTM, sonicated again under the same conditions, and counted using C-Chip counting chambers (Carl Roth, PK36.1).

A total of 1.5 million nuclei were centrifuged as described above, and the pellet was resuspended in 2.6 mL of NEBNext Ultra II Non-Directional RNA Second Strand Synthesis Buffer. Four μL of this mix was added to each well of six fresh 96-well LoBind plates and three 8-tube PCR strips. At this point, three plates were covered with foil seals and stored at −70°C. To each well of the remaining three plates, 1 μL of NEBNext Ultra II Non-Directional RNA Second Strand Synthesis mix containing the second-strand synthesis enzyme was added, and the plates were incubated at 16°C for 2.5 hours, followed by 4°C overnight. DNA Binding Buffer (5 μL; D4004-1-L) was then added to each well. The plates were sealed, vortexed to mix, and incubated at room temperature for 60 minutes. SPRIselect beads (15 μL, 1.5×; Beckman Coulter, B23318) were added to each well, mixed thoroughly by pipetting, and incubated at room temperature for 10 minutes. The beads were cleaned up for each plate using a 96-well magnetic rack and washed twice with 50 μL of freshly prepared 80% ethanol; this additional bead-cleanup step helped reduce the number of uncorrectable barcodes. After the final wash, 7.6 μL of elution buffer (Qiagen, 19086) was added to each well.

One of the three 8-tube PCR strips was used to test different dilutions of the Tn5 transposase. Tn5 (EMBL PepCore), pre-loaded with the N7 oligo (Eurofins), was initially diluted to 0.05 mg/mL and added to TD buffer (20 mM Tris-HCl, pH 7.6; 10 mM MgCl_₂_; 20% dimethylformamide) to generate a series of four 1:2 dilutions. Equal volumes of each Tn5 dilution and nuclei (6.6 μL each) were mixed and incubated at 37°C (lid temperature 47°C) for 15 minutes for tagmentation. The reaction was stopped by adding a transposase stop buffer (0.5% SDS; 10 mg/mL BSA) and incubating at 55°C (lid temperature 65°C) for 15 minutes. The SDS was then quenched by adding 2 μL of 10% Tween-20 to each tube. Next, 2 μL each of indexed TruSeq P5 and indexed Nextera P7 primers (IDT) and 20 μL of 2× NEBNext High-Fidelity PCR Master Mix (NEB, M0541L) were added to each tube, and a 12-cycle PCR was carried out (72°C, 5 minutes; 98°C, 60 seconds; 12 cycles of 98°C for 10 seconds, 63°C for 30 seconds, and 72°C for 60 seconds; 72°C, 5 minutes). Samples were cleaned up with SPRIselect beads (0.8×; Beckman Coulter, B23318) post-PCR. After bead addition, samples were incubated for 10 minutes at room temperature and then placed on a magnetic rack for at least 3 minutes, after which the supernatant was removed. The beads were washed twice as described above and eluted in 11 μL of elution buffer (Qiagen, 19086). Samples were analyzed on a TapeStation 4200 (Agilent), and the optimal Tn5 concentration for the remainder of the experiment was determined based on a library size of 300-700 base pairs. The three main plates were then processed using the chosen Tn5 concentration, with tagmentation, reaction stop, SDS quenching, PCR, and TapeStation analysis performed as above.

The same protocol was used to generate the size and temperature atlas, with modifications to accommodate the scale of the experiment. An additional protease digestion step (2 μL; Qiagen, 19157) was introduced prior to the addition of DNA Binding Buffer to facilitate nuclei breakdown.

### Sequencing, read processing and cell filtering

The size-atlas library was sequenced on an Illumina NovaSeq 6000 (S4 200-cycle kit) using the following sequencing chemistry: Index 1, 10 bp; Index 2, 10 bp; Read 1, 34 bp; Read 2, remaining cycles. Read alignment and gene count matrix generation were performed using the sci-rocket pipeline (https://github.com/lauren-saunders-lab/sci-rocket.git) for sci-RNA-seq3. Reads were aligned to a de novo assembled and annotated transcriptome for *P. morgani*. The sci-rocket pipeline computes a sample-specific threshold for unique molecular identifiers (UMIs) per cell. For downstream filtering, these thresholds were reduced by 25% per sample, with a minimum cutoff of 300 UMIs; cells with UMI counts below this threshold were excluded from further analysis.

The size-temperature atlas library was sequenced on an Element Biosciences AVITI (High 75 bp PE kit) using the following sequencing chemistry: Index 1, 10 bp; Index 2, 10 bp; Read 1, 34 bp; Read 2, remaining cycles. Individual libraries were prepared for each half of the three main plates and sequenced separately across six runs. Sequencing data from the two runs corresponding to each half-plate were combined, and read alignment, gene count matrix generation, and UMI filtering (minimum cutoff 100) were performed for individual plates as described above. Gene count matrices were combined per plate following UMI filtering. An additional doublet-filtering step was performed using scDblFinder (https://github.com/plger/scDblFinder.git) with an expected doublet rate of 10%. Finally, all cells expressing fewer than 75 genes were excluded from further analysis.

### Count matrix pre-processing

After filtering, the size-atlas was processed using Monocle3 (v.1.3.7) workflow defaults unless specified: *estimate_size_factors(), detect_genes(), preprocess_cds(num_dim = 50), align_cds(residual_model_formula_str = “∼log.n.umi + bg_loading”)*, *reduce_dimension(max_components = 2, min_dist= 0.2, umap.n_neighbors = 40L, pre-process_method = “Aligned”)* and finally, *cluster_cells(resolution = 2e-5)*. Clusters predominantly composed of cells originating from only a small number of worms were identified as doublets and excluded from further analysis. The size and temperature atlas was processed with identical parameters, except for the *cluster_cells* parameter *(resolution = 1e-5)*.

### Cell annotation and doublet removal

Cell-type identity for the size-atlas was initially inferred for the 55 global UMAP clusters based on a combination of marker gene expression per cluster and previously published planarian single-cell datasets (*22*, *23*). Next, these clusters were grouped into 14 broad cell types. Each broad cell type was then reprocessed separately to enable higher-resolution annotation and to identify and remove clusters enriched for doublets. Reprocessing was performed using a similar Monocle3 (v.1.3.7) workflow as before: *estimate_size_factors(), detect_genes(), preprocess_cds(num_dim =20), align_cds(residual_model_formula_str = “∼log.n.umi + bg_loading”)*, *reduce_dimension(max_components = 2, min_dist= 0.2, umap.n_neighbors = 40L, pre-process_method = “Aligned”)* and finally, *cluster_cells(resolution = 3e-5)*. Depending on the number of cells the cluster resolution was adjusted. The clustering resolution was adjusted as needed depending on the number of cells in each subset. This reprocessing and doublet-removal procedure was performed twice, resulting in a final annotation of 101 sub-cell types. Lastly, each of the 55 global UMAP clusters was assigned an individual cell type label based on the consensus of the sub-cell type. If no consensus was found at the sub-cell type level, a consensus label based on the broad cell type was used.

### Cell annotation by projection of query data to reference

Cell-type annotations for the query size-temperature atlas were assigned by label transfer from the reference size atlas. The query dataset was subset by temperature, and each subset was independently integrated with the reference and processed using the same Monocle3 workflow and parameters described for the reference above. Each integrated query-reference object was then used for label transfer of both broad cell type and cell type annotations using a k-nearest neighbours (kNN) index, with labels assigned by majority vote of the reference neighbours (k = 20). Cells that did not receive a label during the transfer were excluded from further analysis.

### Cell composition matrix generation and trajectory inference

Cell composition matrix for the size atlas was generated by aggregating single-cell annotations into sample-level counts per cell type. Cell types with fewer than 50 or more than 400,000 cells across the dataset were filtered. In addition, *Cathepsin^+^ transient state cells* were excluded due to their overrepresentation in a single worm. To account for differences in cell recovery per worm, the cell composition matrix was normalized using size factors computed using *estimate_size_factors()*. PCA was then performed on the normalized matrix to visualise variation in cell composition across samples. Trajectory structure was then inferred by learning a principal graph on the PCA embedding using the *learn_graph()* function with parameters set to *ncenter = 7, prune_graph = FALSE* and *minimal_branch_len = 1.* To generate a size-corrected PCA space, we performed ordinary least squares regression for each principal component on body size using the ‘residual_model_formula_str = ∼Lengthmm’ in the align_cds() function in Monocle3 fed with the cell count x worm matrix.

### Covariance analysis

Cell-type covariance across samples was modeled using a Poisson-lognormal framework from PLNmodels (*31*). To control for differences in cell recovery per sample, the logarithm of total cell count per sample was calculated and included as an offset term in all models. The global covariance structure was quantified using a null PLN model. To model size-dependent effects, a second PLN model was fitted with body size as a covariate. This model was fitted with the same offset term as before. Body size was modelled using a natural spline with three degrees of freedom: *model.matrix(∼ ns(size_var, df = 3))*.

### Trajectory clustering

All worms on the trajectory were assigned to a common early-state label (Prebranch_1). A size threshold was defined at the 33rd percentile of body size and worms exceeding this threshold were considered post-branch and partitioned into two groups by k-means clustering (*56*) using *kmeans()* in the PCA space (k=2, seed=123) yielding Branch_1 and Branch_2. Worms assigned to Branch_1 were again clustered in the PCA space (k=2) to resolve Prebranch_2.

### Jensen-Shannon distance analysis

To quantify transcriptomic divergence/similarity between cell types, we pseudobulked the expression profiles by aggregating raw counts per cell type using the *aggregate_gene_expression()* function from Monocle3. Genes with zero total expression across all cell types were removed prior to further analysis. Each cell type expression profile was converted to a probability distribution by dividing gene-level counts by the total count sum per cell type. Jensen-Shannon distances (JSD) were computed between all cell types using the philentropy package in R (https://github.com/drostlab/philentropy.git). JSD values between Neoblast 5 and every other cell type were extracted and ranked.

### Differential cell abundance analysis

To test changes in cell type compositions with temperature and after accounting for body size, we modelled cell type abundances using the Hooke package in R (https://github.com/cole-trapnell-lab/hooke.git). A cell count set was generated using the *new_cell_count_set()* function grouping cells by sample. A model was then fit using the *new_cell_count_model()* function with temperature as the main predictor and body size (as a natural spline with 3 degrees of freedom) included as a nuisance covariate to account size-related variation with parameters set to *vhat_method = “bootstrap”* and num_*bootstraps = 100*. Predicted cell type abundances were estimated at the two temperature extremes (10°C and 22.5°C) with body size held at its mean value. Differential abundance between the two extremes was assessed using *compare_abundances()* function. Cell types were ranked by their q-value to identify those most significantly associated with temperature after accounting for body size.

### Differential gene expression analysis

Differential gene expression analysis across temperatures was performed individually for each cell type using the *fit_models()* function in Monocle3. The model formula included additive terms for temperature (natural spline with 2 degrees of freedom), body size (natural spline with 3 degrees of freedom) and the logarithm of total cell count per sample as an offset term to control for differences in cell recovery per sample. Temperature terms were extracted from the model output, and genes with significant q-values (q<0.05) were filtered to analyze temperature-dependent gene expression changes globally.

---

**Fig. S1.**
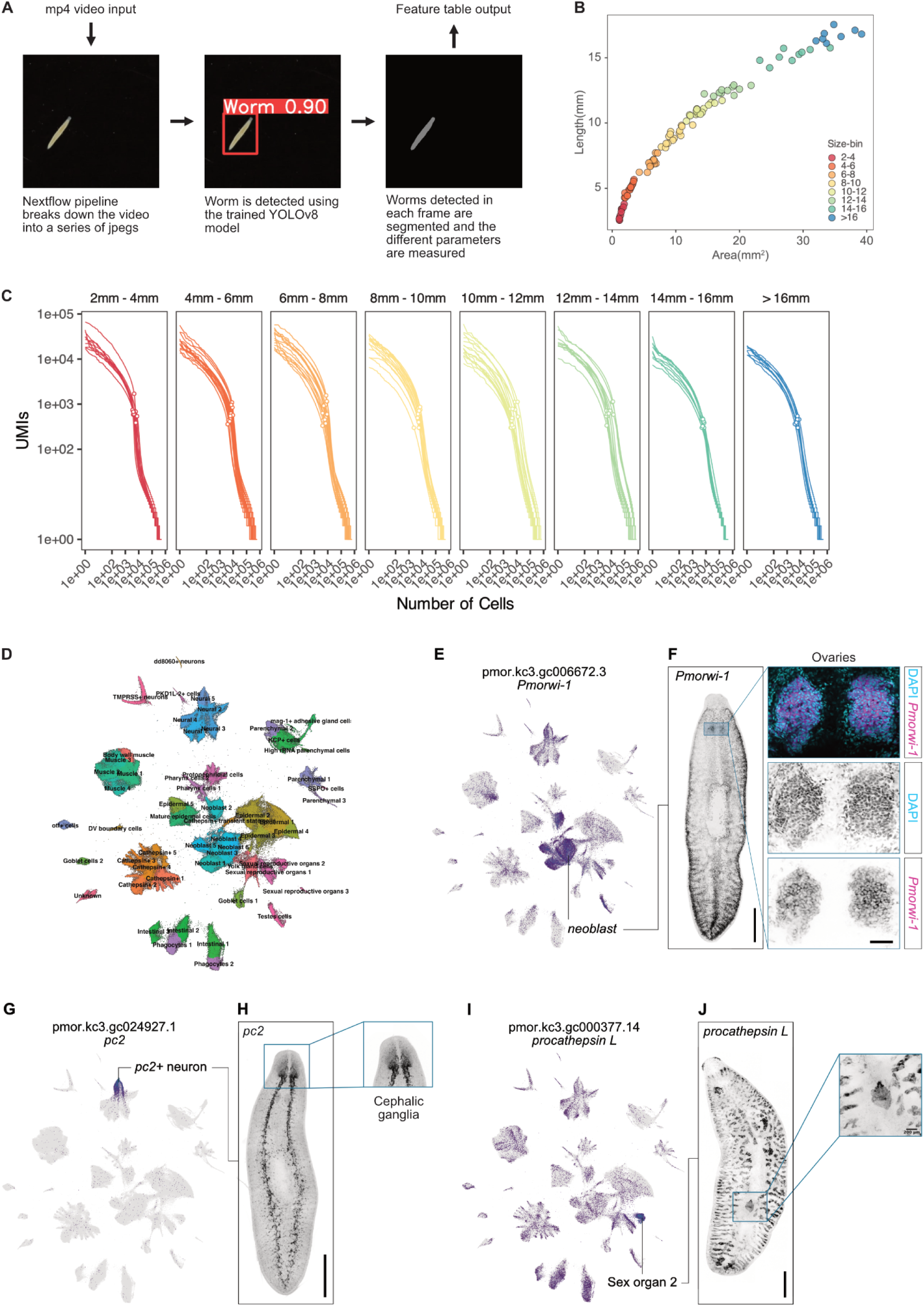
An individually resolved single-cell atlas of *P. morgani* across body size. (**A**) Image-analysis pipeline for measuring body size. A Nextflow pipeline splits each MP4 recording into a series of JPEG frames; worms are detected in each frame with a trained YOLOv8 model (confidence score shown), then segmented to extract morphological features (length, area), which are exported as a per-worm feature table. (**B**) Body length versus area for all worms in the dataset, with each point colored by its assigned size bin (2-4 mm to >16 mm). (**C**) Knee plots of UMI count per cell against ranked cell number, shown per worm and faceted by size bin; open circles mark the sample-specific UMI threshold used for cell calling (adjusted as described in Materials and Methods). (**D**) Two-dimensional UMAP embedding of all profiled cells, colored and labeled by cell type (55 cell types). (**E)** UMAP embedding colored by expression of Pmorwi-1 (pmor.kc3.gc006672.3), a marker of neoblasts. (**F**) Whole-mount FISH of *Pmorwi-1* (left), scale bar: 1mm; insets show a high-magnification view of the ovary with *Pmorwi-1* (magenta) and DAPI (cyan) merged (top), DAPI alone (middle), and *Pmorwi-1* alone (bottom), scale bar: 50 μm. (**G**) UMAP embedding colored by expression of pc2 (pmor.kc3.gc024927.1), a marker of *pc2*+ neurons. (**H**) Whole-mount FISH of *pc2* (left), scale bar: 1mm; inset shows a high-magnification view of the cephalic ganglia. (**I**) UMAP embedding colored by expression of procathepsin L (pmor.kc3.gc000377.14), a marker of sexual reproductive organs 2. (**J**) Whole-mount FISH of *procathepsin L* (left), scale bar: 1mm; inset shows a high-magnification view of sexual reproductive organs 2, scale bar: 200 μm.

**Fig. S2.**
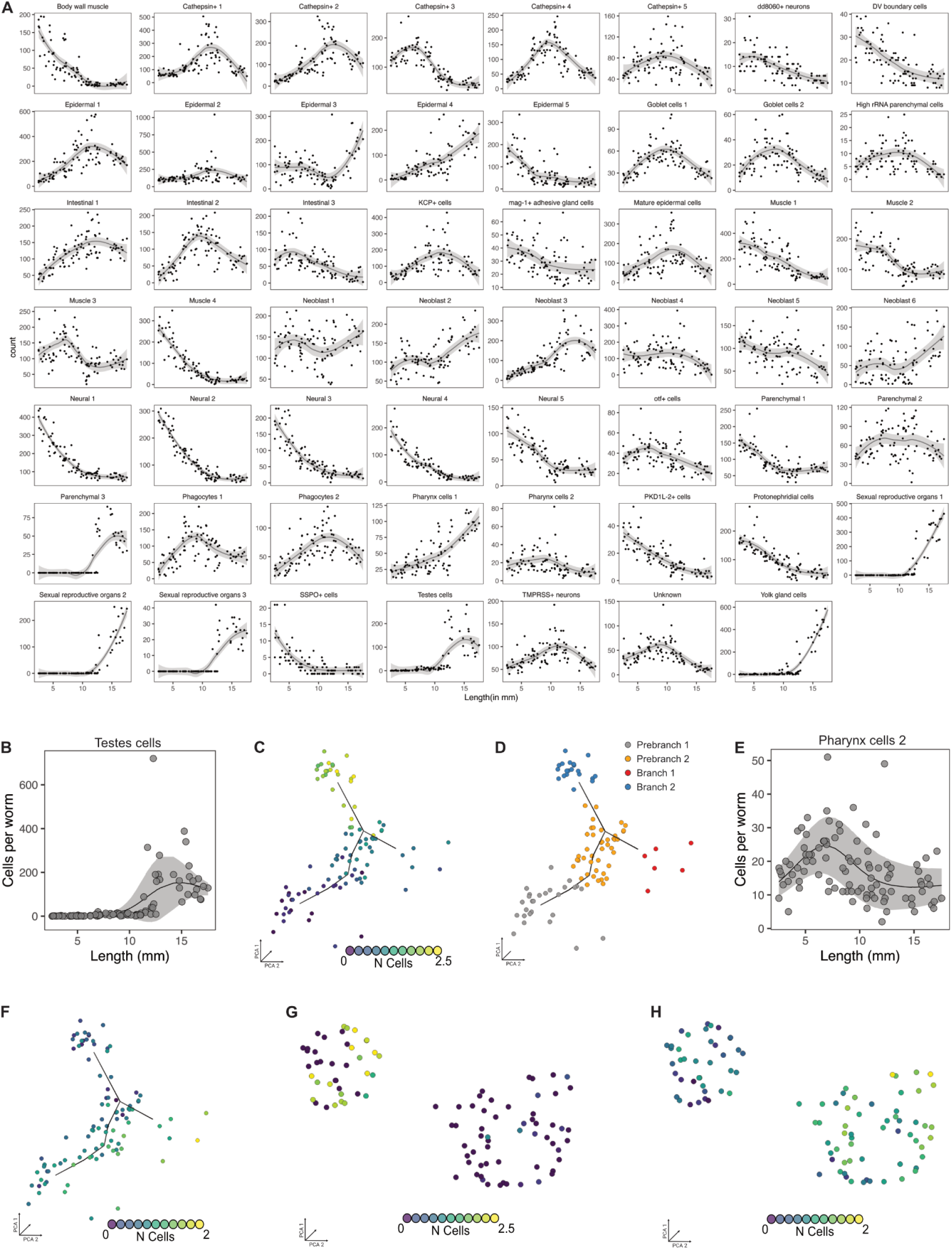
Cell composition dynamics underlying the growth trajectory. (**A**) Allometric scaling of all cell types: cells per worm versus body length for each annotated cell type, with the fitted trend (line) and standard error of the fit (shaded band). (**B**) Sexual reproductive organs 1 abundance (cells per worm) across the sampled size range; line is the fitted trend and the ribbon the standard deviation. (**C**) The same cell type plotted along the growth trajectory on PCA space, with each worm colored by its normalized cell count (log₁₀). (**D**) The growth trajectory is partitioned into pre-branch and post-branch segments (Prebranch 1, Prebranch 2, Branch 1, Branch 2), defined by k-means algorithm. (**E**) Pharynx cells 2 abundance (cells per worm) across the size range, plotted as in (B). (**F**) Pharynx cells 2 along the growth trajectory, colored as in (C). (**G**) Relative abundance of Sexual reproductive organs 2 in size-corrected PCA space, with worms colored by normalized cell count (log₁₀). (**H**) Relative abundance of high rRNA parenchymal cells in size-corrected PCA space, colored as in (G).

**Fig. S3.**
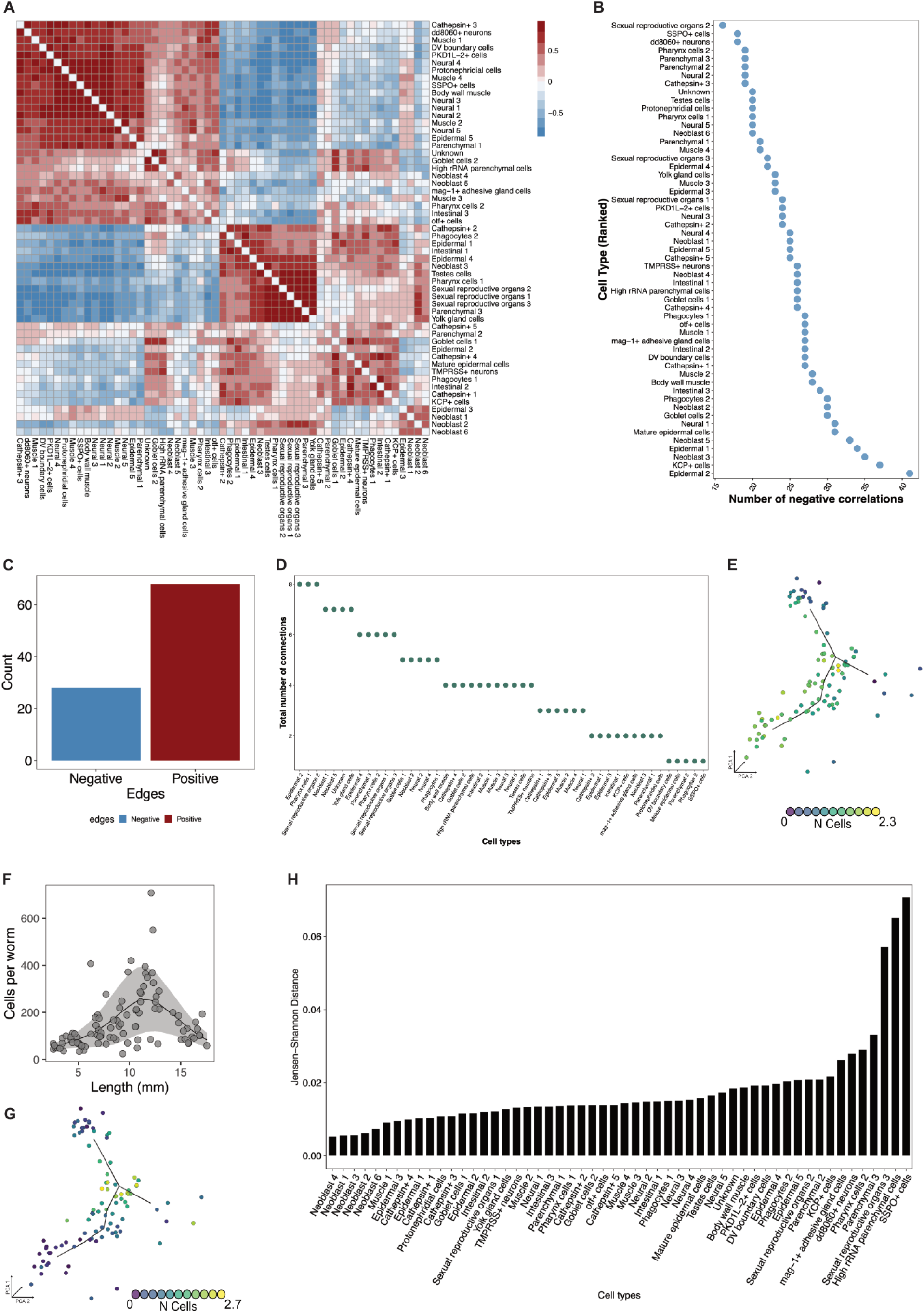
Covariance structure and transcriptional relationships of the cellular drivers. (**A**) Total (uncorrected) cell-cell correlation matrix shown as a heatmap; color indicates correlation strength and sign. (**B**) Cell types ranked by their total number of negative correlations, including both significant and insignificant connections. (**C**) Total counts of significant negative and positive edges across the network. (**D**) Total number of significant connections (edges) per cell type. (**E**) Neoblast 5 abundance along the growth trajectory on PCA space, with each worm colored by normalized cell count (log₁₀). (**F**) *Cathepsin*⁺ 1 abundance (cells per worm) across the sampled size range; line is the fitted trend and the ribbon the standard deviation. (**G**) *Cathepsin*⁺ 1 along the growth trajectory, colored as in (E). (H) Jensen-Shannon distance between the expression profile of Neoblast 5 and every other cell type, ranked from most to least transcriptionally similar; smaller distances indicate closer transcriptional similarity.

**Fig. S4.**
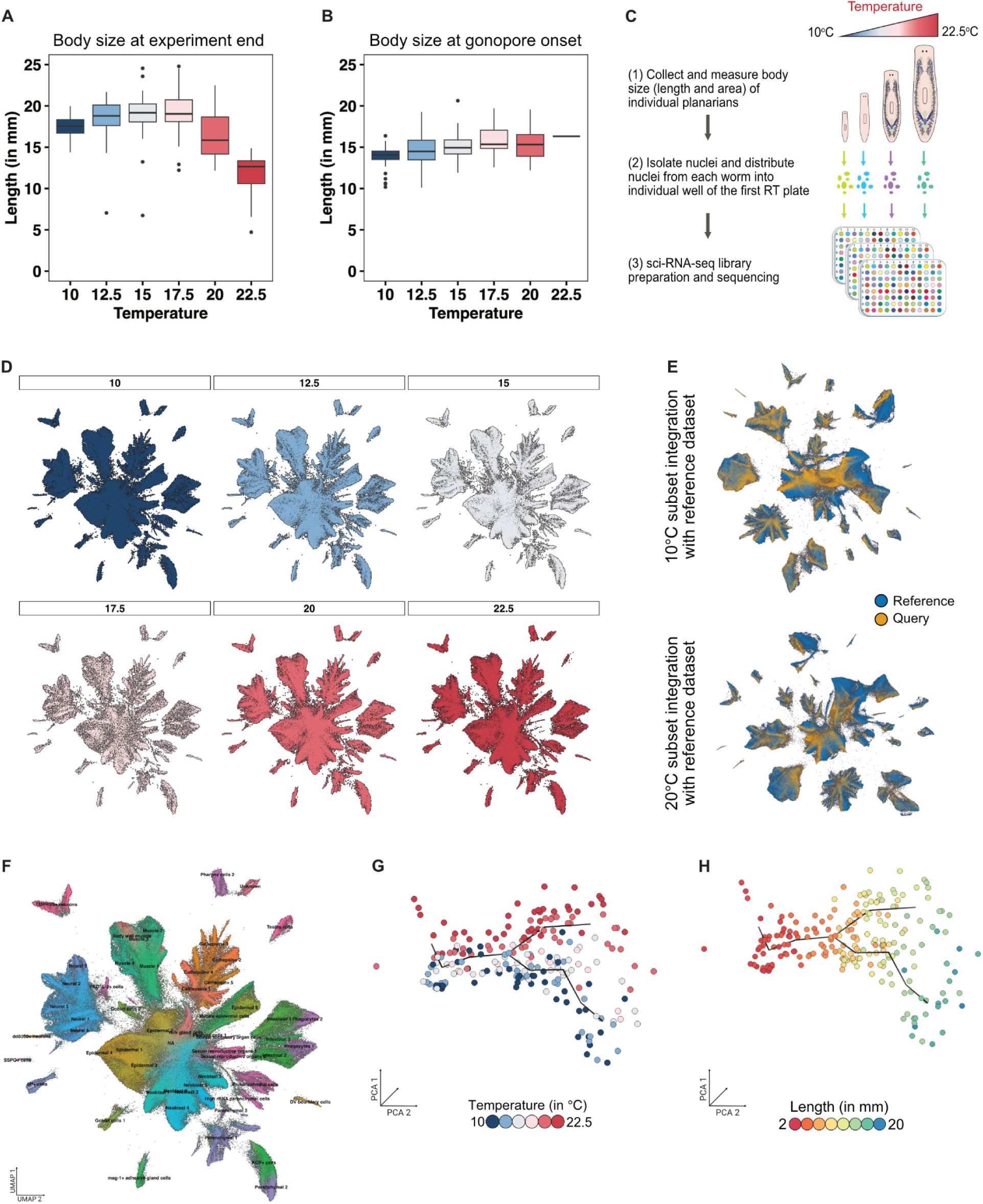
Single-cell profiling of P. morgani across the temperature gradient. (**A**) Body length at the end of the growth experiment across temperature conditions (10-22.5°C); boxes show the median and interquartile range, whiskers the range, and points the outliers. (**B**) Body length at gonopore onset across temperatures, plotted as in (A). (**C**) Workflow for generating the size-temperature atlas: individual planarians grown across the temperature gradient are imaged for body size, then dissociated and distributed one worm per well into the first RT plate for sci-RNA-seq library preparation and sequencing. (**D**) Two-dimensional UMAP embedding of the size–temperature atlas, faceted and colored by temperature condition. (**E**) Co-embedding of each temperature query subset with the reference size atlas, colored by dataset of origin (reference, blue; query, orange); the 10°C and 20°C subsets are shown as examples. (**F**) UMAP embedding colored and labeled by cell type, annotated by label transfer from the reference atlas. (**G**) Growth trajectory inferred for the temperature atlas on PCA space (Monocle3), with each worm colored by temperature treatment. (**H**) The same trajectory colored by body length, showing that the two trajectory branches separate worms of comparable size.

**Fig. S5.**
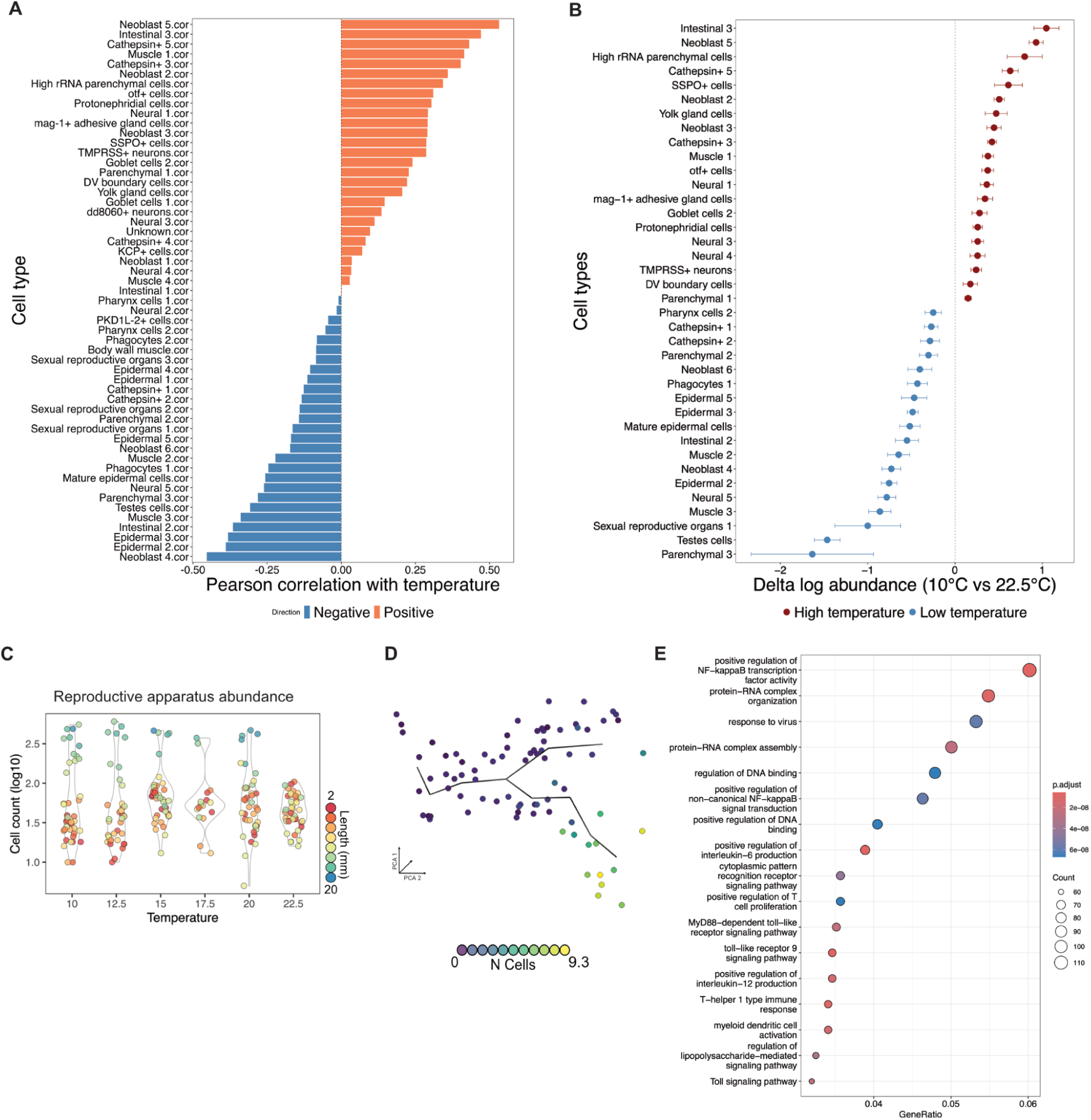
Temperature-dependent changes in cell composition and gene expression. (**A**) Cell types ranked by the Pearson correlation of their abundance with temperature, colored by direction (positive, orange; negative, blue). (**B**) Size-corrected differential abundance between 10°C and 22.5°C (delta log abundance) using Hooke (*32*), with body size included as a covariate; points show the estimate and bars the confidence interval, colored by the temperature at which each cell type is enriched (high, red; low, blue). (**C**) Reproductive-apparatus abundance (cell count, log₁₀) across the temperature gradient; each point is a single worm colored by body length. (**D**) Reproductive-apparatus abundance along the growth trajectory at different temperature on PCA space, with each worm colored by normalized cell count (log₁₀). (**E**) Gene Ontology enrichment for the Neoblast 5 cold-responsive genes, with terms ranked by gene ratio; point size reflects gene count and color the adjusted p-value.

**Table S1.** Top 25 markers per cell type for size-atlas annotation.

**Table S2.** Cell type annotation per cluster used to generate the cell composition matrix.

**Table S3.** Hierarchical annotation for the size-atlas.

**Table S4.** Marker cell types for each trajectory cluster.

**Table S5.** Primer, transcript sequences and transcript IDs used for in situ hybridization

